# Multi-Level Multi-Growth Models: New opportunities for addressing developmental theory using longitudinal designs

**DOI:** 10.1101/2020.10.21.349274

**Authors:** Ethan M. McCormick

**Affiliations:** Department of Psychology and Neuroscience, University of North Carolina, Chapel Hill, North Carolina, 27599

## Abstract

Longitudinal models have become increasingly popular in recent years because of their power to test theoretically derived hypotheses by modeling within-person processes with repeated measures. Growth models constitute a flexible framework for modeling a range of complex trajectories across time in outcomes of interest, including non-linearities and time-varying covariates. However, these models have not thus far been expanded to include the effects of multiple growth processes at once on a single outcome. Here, I outline such an extension, showing how multiple growth processes can be modeled as a specific case of the general ability to include time-varying covariates in growth models. I show that this extension of growth models cannot be accomplished by statistical models alone, and that study design plays a crucial role in allowing for proper parameter recovery. I demonstrate these principles through simulations to mimic important theoretical conditions where modeling the effects of multiple growth processes can address developmental theory including, disaggregating the effects of age and practice or treatment in repeated assessments and modeling age- and puberty-related effects during adolescence. I compare how these models behave in two common longitudinal designs, cohort-sequential and accelerated, and how planned missingness in observations is key to parameter recovery. I conclude with directions for future substantive research using the method outlined here.

## Introduction

An increasing number of fields are beginning to understand the power of longitudinal designs and associated modeling approaches for understanding a broad range of human behavior across time. The developmental sciences are interested in characterizing the maturation (or decline) of cognition and behavior across the lifespan, economists track investment behavior as people age in and out of the workforce, and demographers attempt to model changing populations across decades. However, one major limitation in many models is the ability to model the effects of correlated processes that unfold across time together. For instance, do individuals invest more as they get older or are investment conditions improving across time? The issue of correlated predictors is particularly problematic in the developmental and educational sciences, where age, maturation, and education can all introduce confounds that introduce challenges for both model estimation and inference. For instance, there has been broad interest in how the ability to deploy cognitive control in the service of regulated, flexible behavior across the first two decades of life (Luna, Padmanabhan, & O’Hearn, 2010; Somerville & Casey, 2010). Improvements observed across age have lead naturally to the intuition that these gains in behavioral performance are driven by neurobiological changes associated with maturation. However, this intuition overlooks an important additional force that is exerting influence on neurobiological and behavioral systems, experience. In addition to being biologically more mature, older individuals generally have greater degrees of experience in deploying behavioral strategies across a wide range of contexts. Thus, some of the improvements observed in complex cognitive abilities during adolescence might be due to the effects of everyday practice across long periods of time, even in the absence of biological maturation per se (e.g., in training studies; Braver, Paxton, Locke, & Barch, 2009). However, because this experience plays out along a similar temporal dimension as biological maturation, traditional developmental models are ill-suited to disentangling the disparate forces of development and experience. Despite the many methodological advances in modeling developmental trajectories, new models alone cannot address every theoretical issue without appropriate sampling strategies. Here, I demonstrate how traditional developmental models fail to account for experience when estimating developmental trajectories, and how study design can help address these limitations when combined with a novel extension of traditional models. In particular, I highlight common scenarios where these methods are of interest for addressing long-standing questions for developmental theory.

The aim of the developmental sciences is to understand the course, cause, and consequence of change across time (Curran, Obeidat, & Losardo, 2010). The earliest attempts to understand developmental processes utilized cross-sectional designs (Figure 1A), and this approach remains prevalent in the literature, largely for practical reasons. Cross-sectional designs can either involve comparing differently-aged groups (e.g., developmental stages) or sampling ages from a distribution continuously. However, there are known limitations of these designs stemming from the purely between-person nature of estimated effects, including the inability to test casual relationships or to understand individual differences in trajectories of change (Kraemer, Yesavage, Taylor, & Kupfer, 2000; Louis, Robins, Dockery, Spiro, & Ware, 1986; Maxwell & Cole, 2007). Given these limitations, the developmental sciences have invested heavily in longitudinal (i.e., repeated-measures) designs (Card & Little, 2007) where the same individuals are observed across multiple occasions. Compared with cross-sectional studies, longitudinal designs have increased power to detect developmental effects, allow individuals to be compared to their own baseline, and explicitly model within- and between-person variability (Kraemer et al., 2000; Louis et al., 1986). A common longitudinal design is the cohort-sequential design (Figure 1B), where individuals are measured at the same initial age and followed roughly at equal spacing across time. Many of the largest government-funded longitudinal studies follow this design including the National Longitudinal Surveys (NLS), the UK Biobank, NICHD Study of Early Child Care and Youth Development (SECCYD) Series, and the Adolescent Brain Cognitive Development (ABCD) Study, highlighting the importance of these designs for lifespan developmental research. However, despite the many advantages of these studies, they too face some limitations (aside from the practical challenges of cost and attrition), particularly with respect to disentangling experience and development.

**Figure 1.**
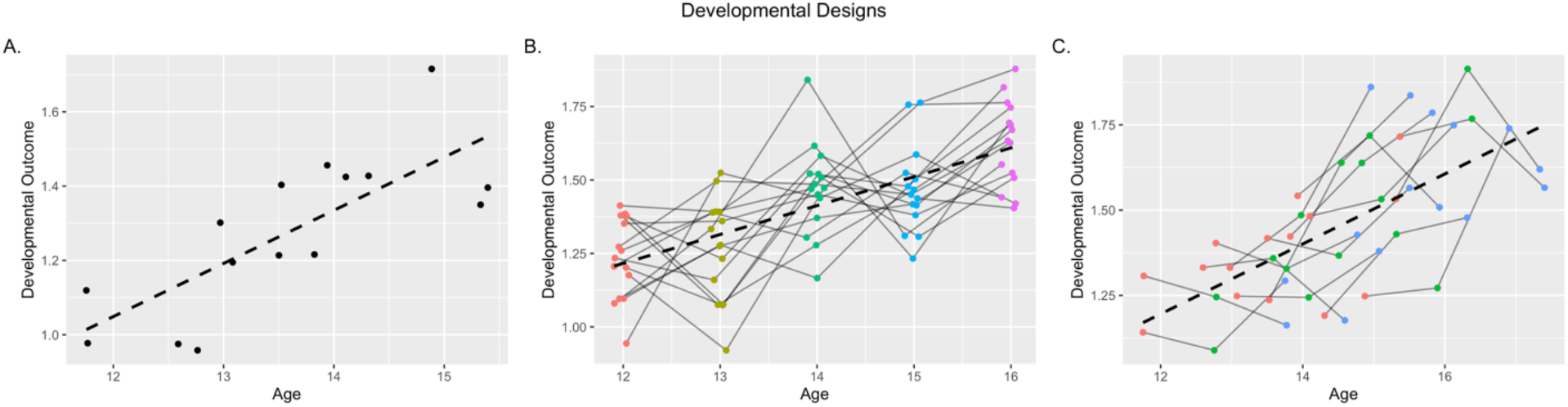
Examples of A) Cross-Sectional, B) Cohort-Sequential, and C) Accelerated Longitudinal developmental designs. While cross-sectional designs rely on one measurement per subject and rely on between-person change to estimate developmental effects (dashed line), cohort-sequential longitudinal designs relay on within-person changes across repeated measurements. Accelerated longitudinal designs combine these approaches by using both between- and within-person change to estimate developmental effects.

The fundamental limitation of cohort-sequential designs is the near or total confounding of age and number of observations (Cook & Ware, 1983; Telzer et al., 2018). This issue has been discussed for over half a century (Bell, 1953; Palmore, 1978) in methodologically focused work, but relatively little attention has been paid to these issues in substantive work since. This oversight is striking, as failure to quantify the effect of repeated measures (e.g., practice, habituation, etc.) presents an existential threat to the internal validity of developmental inferences made from cohort-sequential longitudinal data. The effects of repeated exposure (e.g., practice effects) to observation could artificially enhance (or entirely account for) developmental effects estimated in longitudinal models, or potentially interfere (e.g., habituation) and mask the effects of maturation. Fortunately, an alternative longitudinal design, the mixed or accelerated longitudinal design (Figure 1C), offers a potential solution. Accelerated longitudinal designs combined features of both cross-sectional and cohort-sequential studies (hence mixed), measuring individuals on several occasions but not at the same ages (Bell, 1953). While motivations for adopting accelerated designs have been mostly focused on the cost-efficiency and greater age coverage in shorter study durations (Galbraith, Bowden, & Mander, 2017), these designs also offer a solution to the age-experience confounding seen in cohort-sequential studies by disentangling the effects of repeated observations (i.e., experience) from maturational changes (i.e., development). For instance Cook and Ware (1983) discuss how these models can help to tease apart time, age, and cohort effects in the same model, and other work explicated how specific mathematical formulations could accomplish this (Van’t Hof, Roede, & Kowalski, 1977). The availability of longitudinal data has prompted the development of many models for repeated measures data, including a variety of auto-regressive (Kessler & Greenberg, 1981) latent curve models, and a class of linear regression models known as multi-level (or mixed-effect) models (MLMs; (Bryk & Raudenbush, 1987; Raudenbush & Bryk, 2002). While under conditions common to many cohort-sequential designs, LCMs and MLMs are mathematically isomorphic (Bauer, 2003; Curran, 2003; MacCallum, Kim, Malarkey, & Kiecolt-Glaser, 2010), MLMs are particularly well-suited to the conditions of accelerated longitudinal studies because of their ability to handle the planned-missingness (i.e., individuals are not observed at all possible ages) inherent in the design (Galbraith et al., 2017; Little & Rhemtulla, 2013; Rhemtulla & Little, 2012). While MLMs are applicable to a range of questions, multilevel growth models specifically take the form where an observed outcome is modeled as a function of time (Curran & Bauer, 2011) in a level-one (i.e., person specific) equation

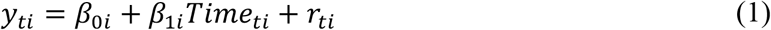

where *y* is the outcome at time *t* for person *i*, *β*_0_ and *β*_1_ are the person-specific intercept and slope (i.e., effect of time) respectively, and *r*_*ti*_ is the time- and person-specific residual. Individuals are allowed to vary by including level-two parameters which model individual variability in intercept and rate of change

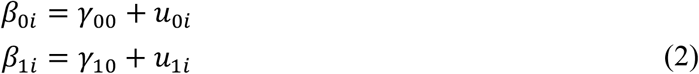

where *γ*_00_ and *γ*_10_ represent the group-level (i.e., fixed effects) intercept and slope, and *μ*_0*i*_ and *μ*_1*i*_ represent individual deviations in those terms (i.e., random effects). By substitution, we can see that the full equation describes a single growth process (across *Time*_*t*_) that allows for estimating individual variability in the initial level and rate of change of *y*_*ti*_.

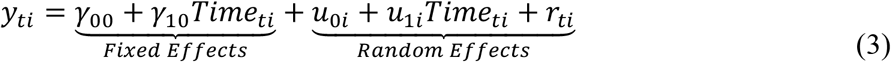

This model could be expanded to include polynomial effects of time, or additional covariates, but this general form constitutes the core of multilevel growth models.

While these models have primarily been used to characterize single growth process (i.e., to model only the effect of time), there are no inherent restrictions on including multiple growth processes in the same model. For instance, modeling two growth processes would involve a simple expansion of Equation 3 to include additional predictors, taking the form:

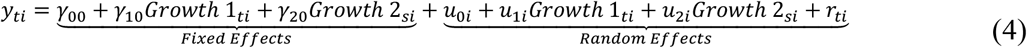

where the growth predictors represent separable constructs of change (e.g., experience and maturation) that unfold across levels of *t* and *s* respectively. This model is not fundamentally different from including any other time-varying covariate, but there are limitations for obtaining proper effect estimates under common conditions. That is, if the two growth processes are highly collinear, there will be instability in effect estimates for either process and standard errors will be inflated (Shieh & Fouladi, 1991), increasing Type II error rates. However, accelerated longitudinal designs offer a potential solution to this limitation since it is possible to attenuate the correlation between different growth processes through the use of planned missingness (Graham, 2009). Combining appropriate sampling designs with the extension to a multilevel, multi-growth modeling framework offers a number of opportunities that can be flexibly employed depending on the specific research question.

In this study, I will highlight the promise of multilevel, multi-growth models (MLMGMs) for addressing developmental theory. To do so, I will develop a series of scenarios where multiple growth processes might be highly entangled. For each scenario, I will highlight how cohort-sequential designs fail to disaggregate between different influences (e.g., age and puberty) on developmental outcomes of interest, and how MLMGMs in combination with accelerated designs address these failures. I show that MLMGMs can be flexibly employed for a range of purposes, from controlling for repeated assessments while estimating developmental effects to probing interactions between chronological age and pubertal maturation. Finally, I outline how sampling strategies can be optimized to take advantage the promise of MLMGMs. While the primary target for this work is the developmental sciences, there are undoubtably applications for these models in other fields where multiple growth processes influence outcomes of interest.

## Methods

### Simulation Design

For each scenario, I simulated data consistent with growth along one or more separable processes (e.g., using variants of Equation 4). For a given scenario, I simulated data to have the structure of both cohort-sequential and accelerated longitudinal designs. I then fit both a properly-specified and mis-specified model (described in each scenario) and demonstrated both how parameters are distributed across different designs and how theoretical inferences would be impacted. To test these questions, I simulated 1000 replicants of each design for each scenario. Data were simulated in R, and all relevant code for replicating data and analyses are available online (https://bit.ly/39Cu7Gw). I plotted results for each scenario in terms of ages that are reasonable for each hypothetical for ease of interpretation and discussion, but they are neither intrinsically meaningful nor meant to directly reflect particular results from real data.

#### Sample Size and Number of Observations

The limitations of common developmental models highlighted in this investigation are not ones that reasonable increases in sample size or number of observations can easily solve. As such, I simulated 750 total observations for each simulated dataset to avoid issues of power and highlight the challenges inherent in confounding age and observations. To mimic real data features, I set 5 observations per person for the cohort-sequential design (n=150), and 3 observations per person for the accelerated data (n=250), with approximately one year between observations (jittered to prevent perfect collinearity *N*∼[0,0.1]). Both of these sample sizes represent reasonably-powered studies for the respective developmental design (Fang, Brooks, Rizzo, Espy, & Barcikowski, 2008).

#### Missing Data

Similar to issues of sample size, I simulated most scenarios to have complete data to focus the scope of the investigation towards issues of theory and inference. This is because planned missing data is accommodated in multilevel models generally and indeed it is what enables their use in accelerated longitudinal data in the first place (Card & Little, 2007). Furthermore, to the extent that missing data influences correlations among growth processes, the most likely contributor to unplanned missing data in longitudinal designs (i.e., attrition) actually attenuates correlations between number of observation and other growth processes. However, one feature of interest to the current investigation is the additional instability that missingness can introduce to parameter estimates in longitudinal data, especially with predictors in the model are highly correlated. To highlight this point, I simulated a separate iteration of one condition in Scenario 1 to have 10% missing data at random (∼ 75 of 750 total observations). The results did not substantively change with the inclusion of missing data, and results are reported in supplemental material (Table S3).

#### Fixed Effects Parameters

Unless otherwise indicated, I simulated regression parameters for the fixed effects to show moderately strong effects (|*γ*_*x*_| = 0.3). This type of effect should be easily detected at the sample sizes shown and clearly highlights the effects of interest in plots. Parameters associated with interactions and polynomials were simulated with weaker absolute effects, and specifics can be found in the description for each scenario below.

#### Random Effects Parameters

Finally, I generated random effects to allow for individual variability in initial level (i.e., intercept, *μ*_0*i*_) and residual variability at level 1 (i.e., *r*_*ti*_). For simplicity, I excluded random effects of slope (i.e., *μ*_0*i*_) from the data generating model; however, MLMGMs, like multilevel, single-growth modes, can be easily expanded to include these effects. Unless otherwise specified, I generated random intercepts (*N*∼[0,0.5]). and residuals (*N*∼[0,1]) from normal distributions. Note that the random effects are expressed as deviations from the fixed effects rather than absolute locations on the scale of the outcome. In later scenarios, I also simulated random slope effects and those effects are described there.

### Parameter Recovery

Observed variables from the data generation step were fit with a set of linear mixed effects models with the lme4 package in R (version 1.1-21; Bates et al., 2015) and significance levels were obtained using the lmerTest package (version 3.1-1) (Kuznetsova, Brockhoff, & Christensen, 2017). I first fit a single growth process MLM (i.e., mis-specified model) as would conventionally be done for the scenario in question (e.g., including only age effects), and then fit an MLMG model with all relevant growth processes included. I extracted both point estimates for the relevant effects, as well as standard errors and significance decisions (i.e., adjusted and unadjusted *p* < or > 0.05). Significance adjustment was done within each model using a Bonferroni correction, and accounted for the number of predictors in a given model (i.e., 1 – 3 predictors depending on the scenario). Based on the point estimate and standard error for each parameter, I also calculated a standardized measure of bias:

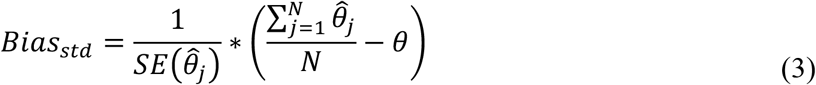

Bias was calculated as the difference between the mean parameter estimate 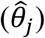 across replications (*j*) and the true generating parameter (*θ*), scaled by the standard error (Collins, Schafer, & Kam, 2001; Gottfredson, Bauer, Baldwin, & Okiishi, 2014). Values represented the distance between the estimated and true parameter as the percentage of the standard deviation of the sampling distribution. For example, a value of .500 would represent a distance of half a standard deviation away from the true parameter. This metric has advantages over significance testing, since differences are very likely to be statistically significant given the large number of iterations, or simple differences that do not take into account the spread of parameter estimates. Previous work has suggested that absolute values above .400 begin to introduce challenges for model performance (Collins et al., 2001). To take a conservative approach, I considered absolute bias values greater than .250 to reflect poor performance. However, no substantive conclusions were altered by the choice of a .250 versus .400 threshold.

## Results

### Scenario Set 1: Age and Practice Effects

Perhaps the most common candidate for a confounding growth process in developmental research is the effect of practice across repeated measures on task measures. At each observation, individuals in the sample are not only older (roughly indicating maturation), but they have been further exposed to the specific task or instrument being used to assess outcomes. Here, I use the example of a difficult cognitive control task. Previous research has shown performance gains across adolescence (Luna et al., 2010; Somerville & Casey, 2010), but these total effects may confound potentially separable impacts of developmental maturation and practice with repeated exposure to the task.

#### Practice Effects Only

Using this framework, I first simulated longitudinal data with no effect of development, but a positive effect of practice (*γ*_*prac*_ = 0.3). In other words, individuals improve their ability to inhibit automatic responses solely through practicing the task, and this practice effect is consistent across age. An exemplar replicant of cohort-sequential and accelerated longitudinal data are plotted below (Figure 2A & B respectively). As expected, the correlation between age and practice was consistently near perfect (*M*_*ρ*_=.998, *SE* < .001) for the cohort design, leading to high levels of variance inflation (*M*_*VIF*_ = 201.58, *SE* = 10.22) and standard errors ~14-times larger than expected. In contrast, these effects were attenuated for the accelerated data with moderate correlations (*M*_*ρ*_=.379, *SE* = .015) and little inflation of standard errors (*M*_*VIF*_ = 1.17, *SE* = .016; standard error inflation = 1.08 times).

**Figure 2.**
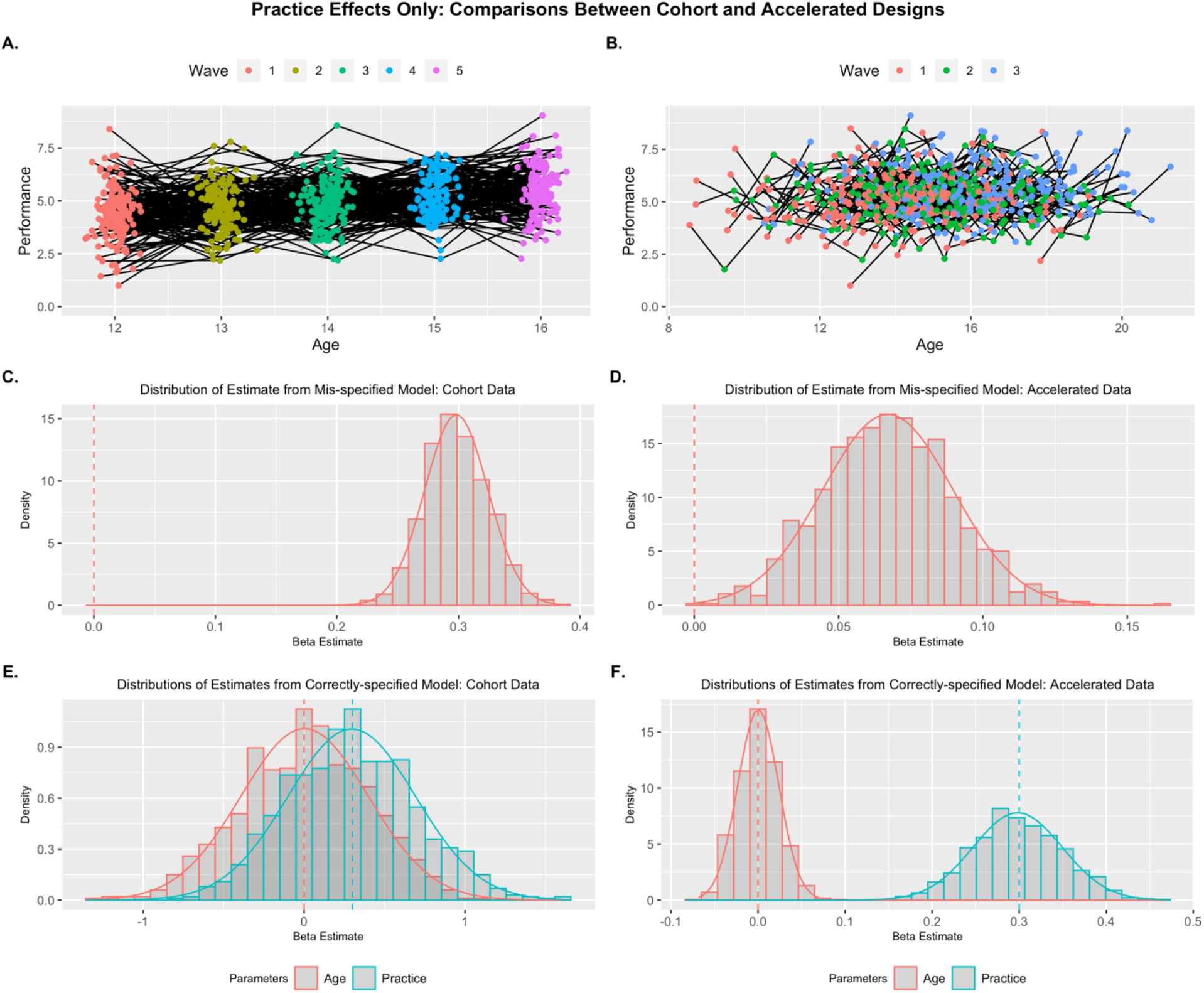
Practice Effects Only: Example replicants for Cohort (A) and Accelerated (B) designs. Distribution of parameter estimates across replicants are presented for mis-specified models where age is the sole predictor (C & D), and properly-specified models where age and practice are included (E & F). Only a MLMG model with accelerated data showed unbiased and precise beta estimates. Generating parameters for each predictor are denoted by vertical dashed lines.

I then fit a traditional, single-growth process model (using age as a predictor) to each dataset and extracted parameter estimates and compared them with their population value (Figure 2C & D). For the cohort data, the practice effect completely aliased as an age effect in the model, whereas there was much less bias in the accelerated data (2.98 vs 11.46; see Table 1 for full parameter details) in the estimate of the age effect due to the attenuated correlation between the two growth predictors. When fitting the correctly-specified multi-growth model, the challenges of disentangling these effects in traditional longitudinal data became clear. While the effects are unbiased (i.e., the distribution center on their respective population values; all bias < .050) even in the presence of high multi-collinearity (Shieh & Fouladi, 1991), the estimates across samples were highly unstable for the cohort data and the distributions of effects for age and practice overlap considerably (Figure 2E). This instability did not lead to inflated false positive rates for the effect of age (age effects were only significant for 4.8% of replicants; 2% when correcting for multiple predictors in the model), but there was a significant elevation of false negatives for the effects of practice, only being significant for 12.6% of replicants without correction (8.2% with correction). This was due to the both increased instability in effect point estimates, as well as inflated standard errors for hypothesis testing. Neither of these issues were present in the accelerated longitudinal data, as there was virtually no bias in either effect estimates and false positive (3.8% for effects of age without correction; 1.5% with correction) and false negative (0% for effects of practice) rates were appropriately low. All models correctly captured the variance of the residual (*r*_*ti*_) and person-level (*τ*_00_) random effects, with some downward bias (Shieh & Fouladi, 1991).

**Table 1.** Estimated Effects for Growth Models from Scenario Set 1: Single Effects.

|  | Cohort-Sequential Design |  |  |  |  |  | Accelerated Design |  |  |  |  |  |
| --- | --- | --- | --- | --- | --- | --- | --- | --- | --- | --- | --- | --- |
|  | Mean Est. | Std. Err. | Min | Max | Std. Bias | Prop. Sig.(adj.) | Mean Est. | Std. Err. | Min | Max | Std. Bias | Prop. Sig.(adj.) |
| *Practice Effects Only* |  |  |  |  |  |  |  |  |  |  |  |  |
| $\rho_{growth}$ | .998 | .000 | .997 | .998 | | | .379 | .015 | .335 | .426 | | |
| *VIF* | 201.6 | 10.2 | 171.7 | 233.8 |  |  | 1.17 | .015 | 1.13 | 1.22 |  |  |
| $\gamma_{age (mis)}$ | .298 | .026 | .223 | .385 | **11.46** | 1 | .067 | .023 | .002 | .162 | **2.98** | .865 |
| $\sigma (mis)$ | 1.00 | .029 | .903 | 1.09 | .011 | | 1.03 | .033 | .920 | 1.15 | **.784** | |
| $\sqrt{\tau_{00} (mis)}$ | .496 | .054 | .303 | .669 | -.082 | | .497 | .062 | .245 | .660 | -.042 | |
| $\gamma_{age}$ | .003 | .395 | -1.31 | 1.12 | .008 | .048 (.020) | .000 | .023 | -.068 | .089 | .014 | .038 (.015) |
| $\gamma_{prac}$ | .296 | .396 | -.777 | 1.60 | -.009 | .126 (.082) | .298 | .051 | .147 | .470 | -.034 | 1 (1) |
| $\sigma$ | 1.00 | .029 | .903 | 1.08 | -.005 | | .999 | .032 | .888 | 1.12 | -.021 | |
| $\sqrt{\tau_{00}}$ | .496 | .054 | .302 | .663 | -.082 | | .498 | .057 | .290 | .648 | -.035 | |
| *Age Effects Only* |  |  |  |  |  |  |  |  |  |  |  |  |
| $\rho_{growth}$ | .998 | .000 | .997 | .998 | | | .378 | .014 | .335 | .417 | | |
| *VIF* | 202.1 | 10.7 | 171.7 | 243.4 |  |  | 1.17 | .015 | 1.13 | 1.21 |  |  |
| $\gamma_{age (mis)}$ | .299 | .025 | .218 | .373 | -.024 | 1 | .301 | .021 | .233 | .361 | .040 | 1 |
| $\sigma (mis)$ | 1.00 | .029 | .893 | 1.10 | .021 | | 1.00 | .031 | .924 | 1.09 | .008 | |
| $\sqrt{\tau_{00} (mis)}$ | .498 | .053 | .314 | .646 | -.031 | | .497 | .058 | .257 | .658 | -.057 | |
| $\gamma_{age}$ | .298 | .387 | -.848 | 1.64 | -.005 | .114 (.071) | .301 | .025 | .223 | .379 | .041 | 1 (1) |
| $\gamma_{prac}$ | .001 | .388 | -1.41 | 1.15 | .004 | .045 (.023) | -.001 | .051 | -.150 | .166 | -.016 | .044 (.020) |
| $\sigma$ | 1.00 | .029 | .893 | 1.10 | .022 | | 1.00 | .031 | .924 | 1.09 | .008 | |
| $\sqrt{\tau_{00}}$ | .498 | .054 | .314 | .646 | -.030 | | .497 | .058 | .257 | .658 | -.057 | |

Note: Mean Est. is the expected parameter value across replicants and the Std. Err is the standard deviation of the sampling distribution. The proportion significant (Prop. Sig) represents the number of inferential tests that return a significant result for each parameter (unadjusted first, adjusted in parentheses). Std. Bias is the standardized bias estimate; values > .25 are bolded and represent unacceptable bias. The proportion significant only includes significant results where the inference is correct (e.g., significant negative effects are not included if the population parameter is positive). Rhos ( $\rho_{growth}$ ) represent the correlation between growth predictors, VIF represents variance inflation factor, gammas ( $\gamma$ ) represent the fixed effects, sigma ( $\sigma$ ) is the square-root of the residual random variance, and tau ( $\tau_{00}$ ) is the person-level random variance (square-root shown for consistency). The (mis) denotes parameters from the mis-specified (i.e., age-only) model.

#### Age Effects Only

I next simulated a set of data to only contain effects of age and not practice to highlight the specificity of MLMGMs. That is when the potential confounding growth process of practice did not contribute to developmental outcomes, MLMGMs do not produce false positives. Here, only positive age effects (*γ*_*age*_ = 0.3) were included, suggesting the behavioral improvements in inhibitory control only impacted solely by maturation. All results were the mirror image of these for the effects of practice only (see Table 1 for complete parameter details; Figure S1), and show that MLMG models are capable of accurately disaggregating the effects of practice and age. While in this condition, the effect in the longitudinal data could be captured using only the single growth effect of age, that could only be ensured in a simulation where the true data-generating model is known; an unlikely assumption for data collected in the wild.

#### Additive Effects of Age and Practice

Combining the previous two simulations, I then simulated data to contain effects of both age and practice (*γ′s* = 0.3). This is a more plausible scenario than the two previous conditions as both maturity and exposure likely contribute to increases in task performance. However, based on results thus far, we would expect that effects in cohort-sequential models with only an age predictor would be inflated relative to the true effect due to the high correlation (*M*_*ρ*_=.998, *SE* < .001) and unstable due to the variance inflation (*M*_*VIF*_ = 202.18, *SE* = 10.67) if both predictors were included. Indeed, in this additive effects model, a single growth model with the effect of age substantially inflates the predicted effect (*Bias*_*std*_ = 11.32, *SE* = .026) in the cohort data, while inflating the effect much less in the accelerated design (*Bias*_*std*_ = .2.91, *SE* = .022; Figure 3C & D). The MLMG models show unbiased effect estimates for both age and practice for both design types, but false negatives are controlled appropriately only in the accelerated data (Figure 3; Table 2).

**Figure 3.**
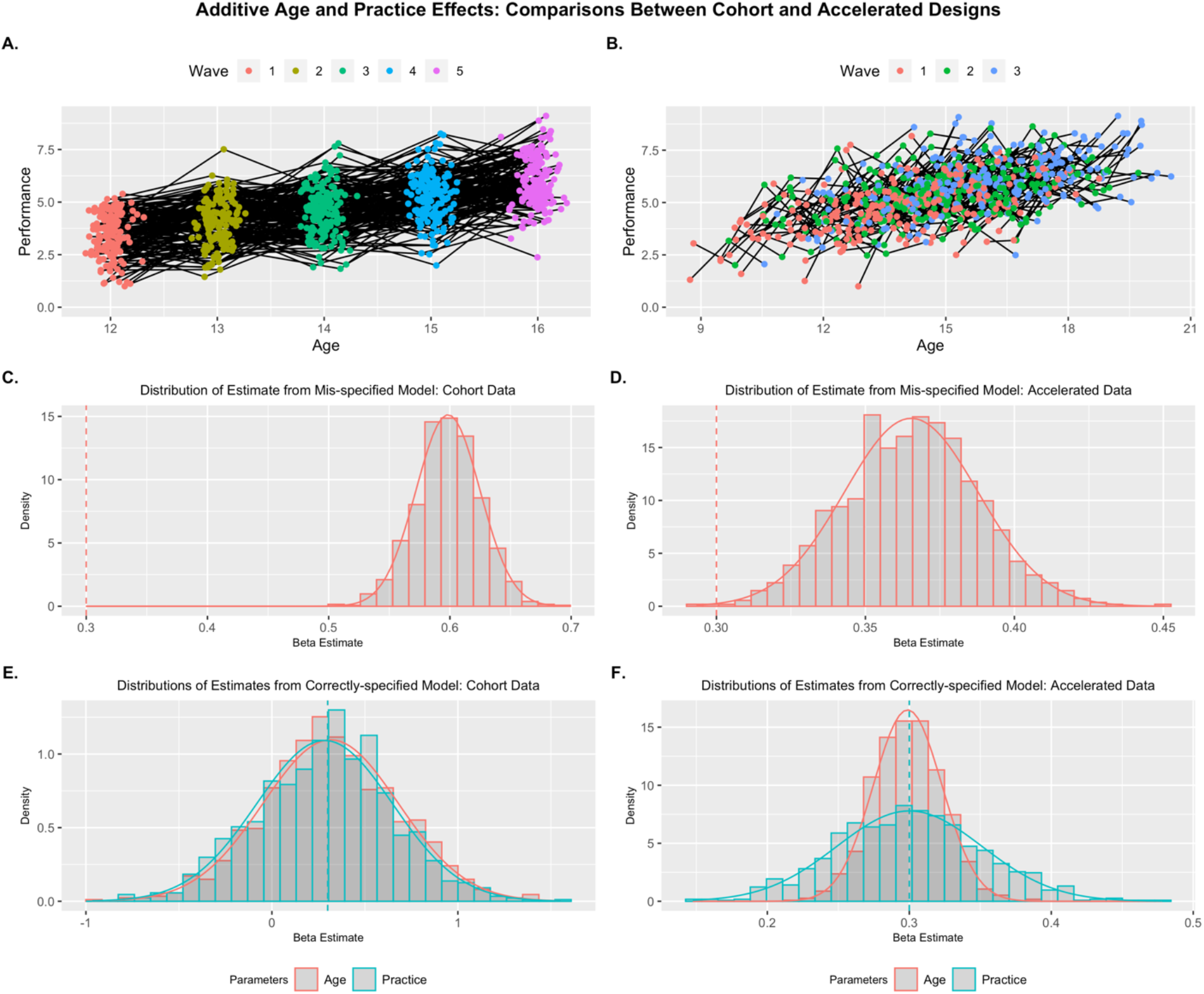
Additive Practice and Age Effects: Example replicants for Cohort (A) and Accelerated (B) designs. Distribution of parameter estimates across replicants are presented for mis-specified models where age is the sole predictor (C & D), and properly-specified models where age and practice are included (E & F). Only a MLMG model with accelerated data showed unbiased and precise beta estimates. Generating parameters for each predictor are denoted by vertical dashed lines.

**Table 2.**
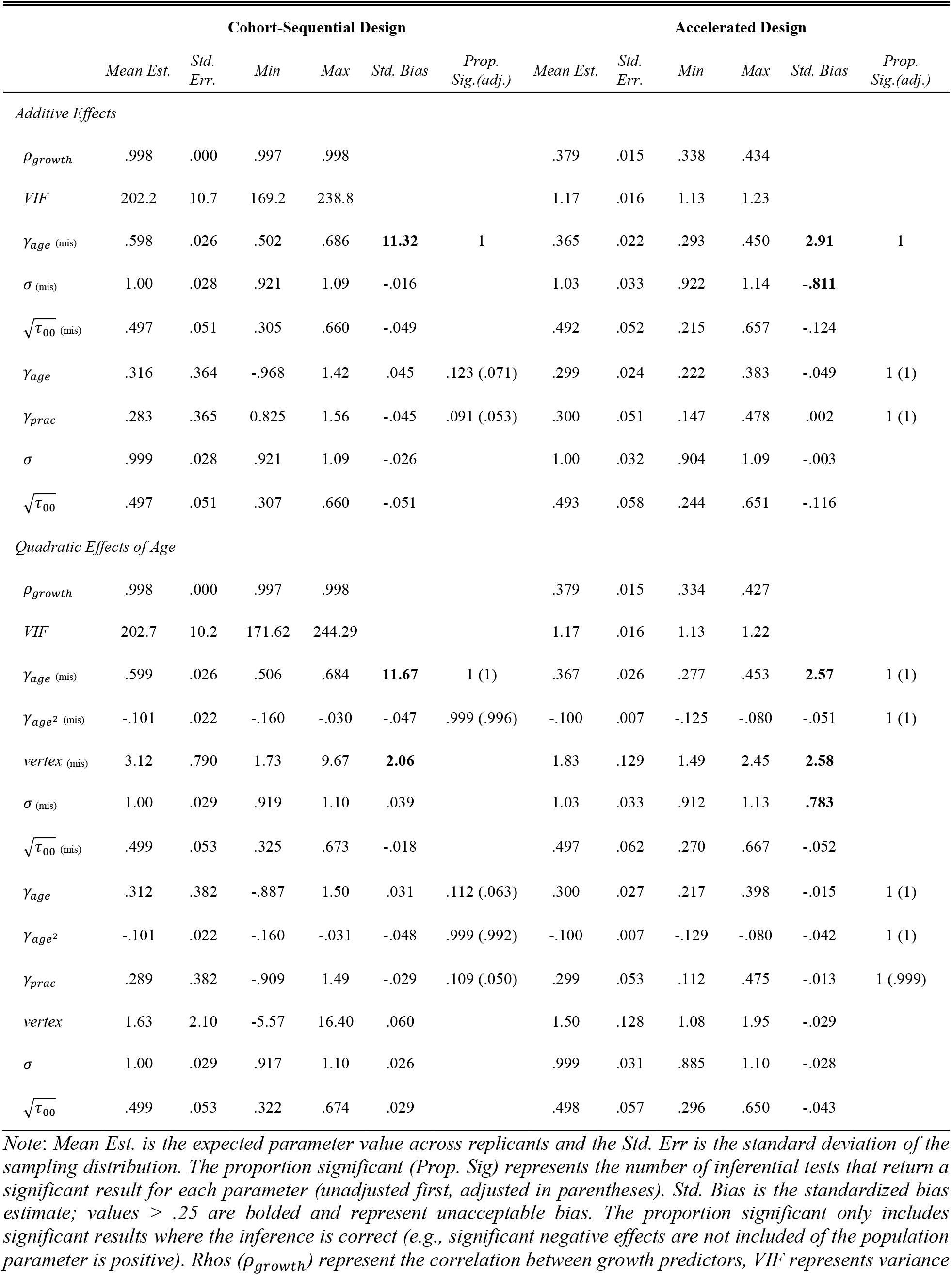
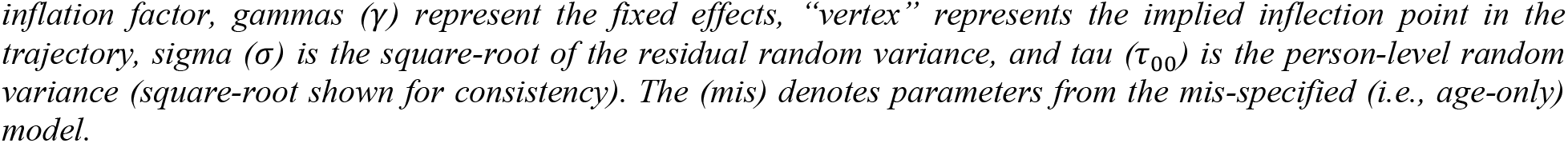
Estimated Effects for Growth Models from Scenario Set 1: Additive Effects.

|  | Cohort-Sequential Design |  |  |  |  |  | Accelerated Design |  |  |  |  |  |
| --- | --- | --- | --- | --- | --- | --- | --- | --- | --- | --- | --- | --- |
|  | Mean Est. | Std. Err. | Min | Max | Std. Bias | Prop. Sig.(adj.) | Mean Est. | Std. Err. | Min | Max | Std. Bias | Prop. Sig.(adj.) |
| *Additive Effects* |  |  |  |  |  |  |  |  |  |  |  |  |
| $\rho_{growth}$ | .998 | .000 | .997 | .998 | | | .379 | .015 | .338 | .434 | | |
| VIF | 202.2 | 10.7 | 169.2 | 238.8 |  |  | 1.17 | .016 | 1.13 | 1.23 |  |  |
| $\gamma_{age} \text{ (mis)}$ | .598 | .026 | .502 | .686 | **11.32** | 1 | .365 | .022 | .293 | .450 | **2.91** | 1 |
| $\sigma \text{ (mis)}$ | 1.00 | .028 | .921 | 1.09 | -.016 | | 1.03 | .033 | .922 | 1.14 | **-.811** | |
| $\sqrt{\tau_{00}} \text{ (mis)}$ | .497 | .051 | .305 | .660 | -.049 | | .492 | .052 | .215 | .657 | -.124 | |
| $\gamma_{age}$ | .316 | .364 | -.968 | 1.42 | .045 | .123 (.071) | .299 | .024 | .222 | .383 | -.049 | 1 (1) |
| $\gamma_{prac}$ | .283 | .365 | 0.825 | 1.56 | -.045 | .091 (.053) | .300 | .051 | .147 | .478 | .002 | 1 (1) |
| $\sigma$ | .999 | .028 | .921 | 1.09 | -.026 | | 1.00 | .032 | .904 | 1.09 | -.003 | |
| $\sqrt{\tau_{00}}$ | .497 | .051 | .307 | .660 | -.051 | | .493 | .058 | .244 | .651 | -.116 | |
| *Quadratic Effects of Age* |  |  |  |  |  |  |  |  |  |  |  |  |
| $\rho_{growth}$ | .998 | .000 | .997 | .998 | | | .379 | .015 | .334 | .427 | | |
| VIF | 202.7 | 10.2 | 171.62 | 244.29 |  |  | 1.17 | .016 | 1.13 | 1.22 |  |  |
| $\gamma_{age} \text{ (mis)}$ | .599 | .026 | .506 | .684 | **11.67** | 1 (1) | .367 | .026 | .277 | .453 | **2.57** | 1 (1) |
| $\gamma_{age^2} \text{ (mis)}$ | -.101 | .022 | -.160 | -.030 | -.047 | .999 (.996) | -.100 | .007 | -.125 | -.080 | -.051 | 1 (1) |
| $vertex \text{ (mis)}$ | 3.12 | .790 | 1.73 | 9.67 | **2.06** | | 1.83 | .129 | 1.49 | 2.45 | **2.58** | |
| $\sigma \text{ (mis)}$ | 1.00 | .029 | .919 | 1.10 | .039 | | 1.03 | .033 | .912 | 1.13 | **.783** | |
| $\sqrt{\tau_{00}} \text{ (mis)}$ | .499 | .053 | .325 | .673 | -.018 | | .497 | .062 | .270 | .667 | -.052 | |
| $\gamma_{age}$ | .312 | .382 | -.887 | 1.50 | .031 | .112 (.063) | .300 | .027 | .217 | .398 | -.015 | 1 (1) |
| $\gamma_{age^2}$ | -.101 | .022 | -.160 | -.031 | -.048 | .999 (.992) | -.100 | .007 | -.129 | -.080 | -.042 | 1 (1) |
| $\gamma_{prac}$ | .289 | .382 | -.909 | 1.49 | -.029 | .109 (.050) | .299 | .053 | .112 | .475 | -.013 | 1 (.999) |
| $vertex$ | 1.63 | 2.10 | -5.57 | 16.40 | .060 | | 1.50 | .128 | 1.08 | 1.95 | -.029 | |
| $\sigma$ | 1.00 | .029 | .917 | 1.10 | .026 | | .999 | .031 | .885 | 1.10 | -.028 | |
| $\sqrt{\tau_{00}}$ | .499 | .053 | .322 | .674 | .029 | | .498 | .057 | .296 | .650 | -.043 | |

Note: Mean Est. is the expected parameter value across replicants and the Std. Err is the standard deviation of the sampling distribution. The proportion significant (Prop. Sig) represents the number of inferential tests that return a significant result for each parameter (unadjusted first, adjusted in parentheses). Std. Bias is the standardized bias estimate; values > .25 are bolded and represent unacceptable bias. The proportion significant only includes significant results where the inference is correct (e.g., significant negative effects are not included of the population parameter is positive). Rhos ( $\rho_{growth}$ ) represent the correlation between growth predictors, VIF represents variance
*inflation factor, gammas ( $\gamma$ ) represent the fixed effects, “vertex” represents the implied inflection point in the trajectory, sigma ( $\sigma$ ) is the square-root of the residual random variance, and tau ( $\tau_{00}$ ) is the person-level random variance (square-root shown for consistency). The (mis) denotes parameters from the mis-specified (i.e., age-only) model.*

#### Shifting Quadratic Effects of Age

While I have only considered linear effects thus far, there are many plausible scenarios involving non-linear effects of age (or other growth predictors). Here the confounding of practice effects can be more pernicious, especially with respect to inference, because of the way higher-order polynomial effects are built. Here I considered a relatively straightforward scenario with a decelerating quadratic effect of age 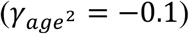 combined with a positive linear effect of age and practice (*γ′s* = 0.3). This pattern of effects might be characteristic of some adolescent-specific advantage in performance, but that adult performance would still exceed childhood levels, combined with a within-person effect of practice boosting performance.

There are two primary inferential issues that could be impacted in this type of scenario. First, as we have seen previously, failure to consider practice effects (or considering them but in cohort-sequential designs) will seriously impair our ability to accurately characterize age-related changes in performance. However, the introduction of the quadratic term provides an added temptation: to draw conclusions about the specific location of the peak of the trend in performance. We know that the specific shape (and indeed order) of the polynomial effect can be influenced by boundary observations at the low or high end of the age distribution. Nevertheless, these peaks (or troughs) are almost irresistible when drawing substantive conclusions about developmental theory. As such, I explored the recovery of specific model parameters similar to earlier scenarios, as well as the implied vertex of the quadratic trend.

As in the scenario with only additive effects of age and practice, there was high collinearity between the linear predictors of age and experience (*M*_*ρ*_=.998, *SE* < .001). This issue did not impact the quadratic predictor of age because the linear predictor was centered before being squared (chapter 2) (Aiken, West, & Reno, 1991). When fitting models with only effects of age (linear and quadratic), the pattern of results was identical to the linear case, where age absorbed the practice effect fully in the cohort data (*Bias*_*std*_ = 11.67, *SE* = .026) and to a lesser degree in the accelerated data (*Bias*_*std*_ = 2.57, *SE* = .026; Figure 4C & D; see Table 2 for full parameter details). In contrast, the beta estimates for the quadratic effect were unbiased and precise (due to the uncorrelated nature of this predictor). However, the implied vertex in the developmental trend was still shifted due to the biased point estimate of the linear predictor (Figure 4E & F; note the difference in x-axis spread for the cohort data). Adopting the proper model in both designs ameliorated the bias in effect estimates, although as before, the model from the cohort data was very imprecise, with large spreads in parameter estimates and implied vertices (Figure 4G & I), as well as a high rate of false negatives (88.8% for age estimate; 89.1% for practice estimate). Of particular concern for interpretation, the vertex for these models ranges across more than a full decade. In contrast, the properly-specified MLMGM with the accelerated design produced unbiased and precise estimates of all parameters (Figure 4H & J) with no false negatives. As expected, all models recovered the quadratic effect successfully with almost no false negatives (0.1 % for cohort data, 0% for accelerated data).

**Figure 4.**
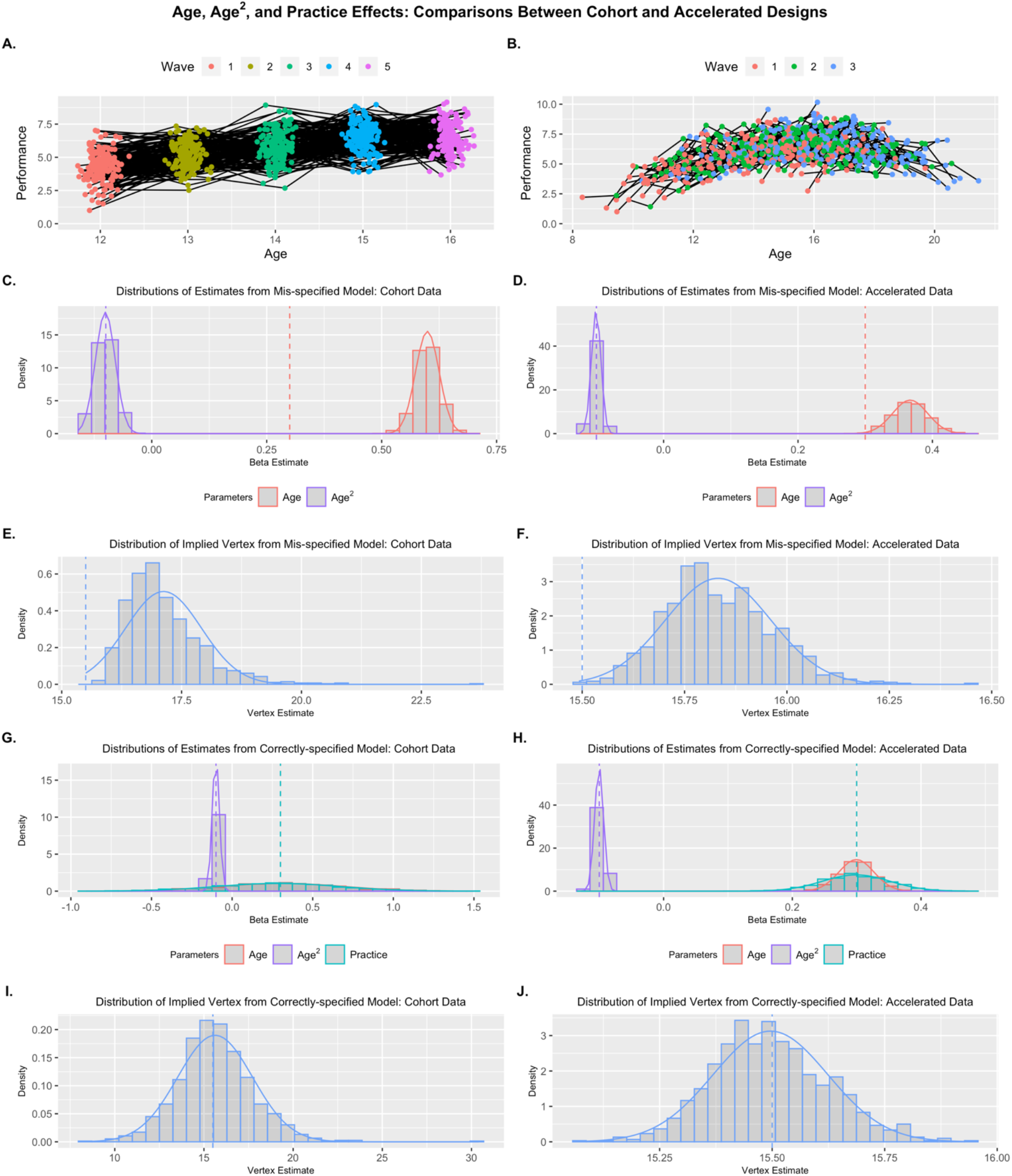
Shifting Quadratic Effects of Age: Example replicants for Cohort (A) and Accelerated (B) designs. Distribution of parameter estimates (C & D) and implied vertices (E & F) across replicants are presented for mis-specified models where age is the sole predictor, and properly-specified models where age and practice are included (G – J). Only a MLMG model with accelerated data showed unbiased and precise beta estimates and implied vertices. Generating parameters for each predictor are denoted by vertical dashed lines.

These results highlight a wrinkle in what the reader might have thought could be a saving grace for estimating MLMG models in cohort data. That is, unlike in the purely linear scenarios considered thus far, practice effects are more often likely to follow a non-linear pattern across time, where changes between the first and second observation are large and then subsequent changes are smaller. Any polynomial effect of practice would follow the same pattern as the quadratic age effect in this scenario, being estimated precisely and without bias. However, to capture this in the model, the linear predictor would need to be included as well, introducing issues with both the linear practice estimate, as well as any linear age effect being modeled. As such, no matter what complex trends the outcome of interest shows across either age or practice, the necessity of including the linear terms in the model poses problems for appropriate inference. Furthermore, this instability in estimates produces additional challenges for interpreting the non-linear terms (despite being properly estimated) since the vertex of the trend depends, in-part, on the linear term and can take substantively different forms (e.g., inverted-U versus a reflected exponential) as a result.

### Scenario Set 2: Interference Effects

Of course, the effects of repeated assessments can go beyond practice, which can generally be assumed to boost task performance. Other confounders may operate against the axis of the primary growth process of interest. In the next simulations, I considered cases when multiple growth processes interfere with one another, using examples common in developmental research.

#### Habituation Effects

An prime example of this kind of interferences is the process of habituation in emotional reactivity to a set of stimuli. This could take the form of a startle response, rating of negative emotions (e.g., anger, disgust), or brain response (e.g., dorsal anterior cingulate or amygdala) that decreases once stimuli are less unexpected (Breiter et al., 1996; Denny et al., 2014; Ellwanger, Geyer, & Braff, 2003). In contrast, we might expect general developmental increases in negative emotionality (e.g., depression) across some periods of development (e.g., adolescence; Cyranowski, Frank, Young, & Shear, 2000; Garber, Keiley, & Martin, 2002), which habituation effects would work to attenuate. To demonstrate this kind of masking by the confounder, I simulated data to have a positive effect of age (*γ*_*age*_ = 0.3) but a negative effect of exposure (*γ*_*exp*_ = −0.3) indicating habituation across repeated measures (Figure). As seen with the additive models, the age-only growth model in the cohort data showed substantial bias (*Bias*_*std*_ = −12.17, *SE* = .025), and results would infer that there is no change in negative reactivity across age (Figure 5). In the age-only model with the accelerated data, the correlation between age and exposure does attenuate the true effect of age somewhat, but much less (*Bias*_*std*_ = −3.02, *SE* = .022) and all replicants showed a significant positive effect of age despite this attenuation (see Table 3 for full details). With the correctly-specified MLMGM model, the effect estimates showed the same instability as previously shown, while in the accelerated design, the two were well-separated. Additionally, the MLMGM with accelerated data provided appropriate false negative control.

**Figure 5.**
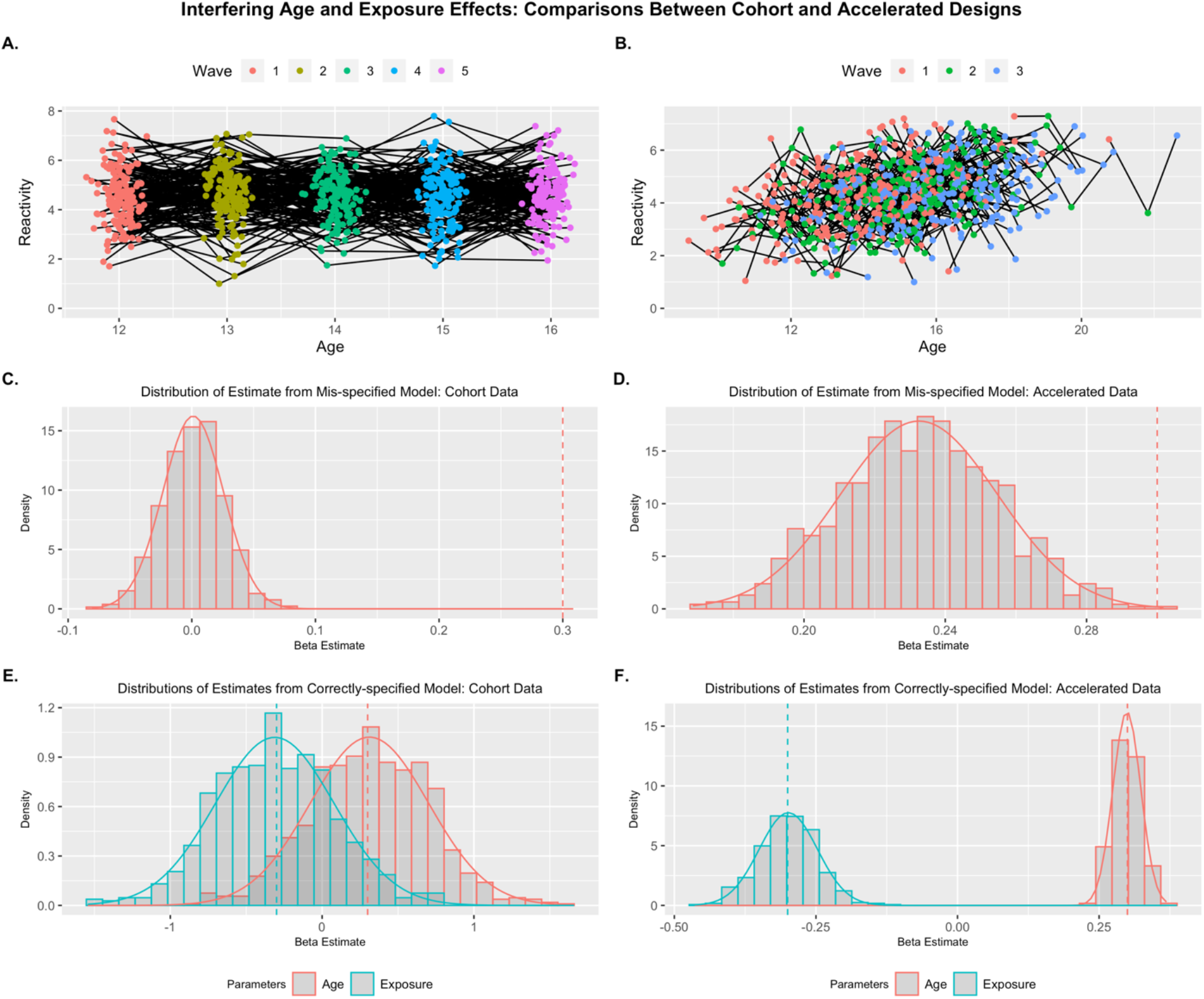
Interfering Age and Exposure Effects: Example replicants for Cohort (A) and Accelerated (B) designs. Distribution of parameter estimates across replicants are presented for mis-specified models where age is the sole predictor (C & D), and properly-specified models where age and exposure are included (E & F). Only a MLMG model with accelerated data showed unbiased and precise beta estimates. Generating parameters for each predictor are denoted by vertical dashed lines.

**Table 3.** Estimated Effects for Growth Models from Scenario Set 2: Interference Effects.

|  | Cohort-Sequential Design |  |  |  |  |  | Accelerated Design |  |  |  |  |  |
| --- | --- | --- | --- | --- | --- | --- | --- | --- | --- | --- | --- | --- |
|  | Mean Est. | Std. Err. | Min | Max | Std. Bias | Prop. Sig. (adj.) | Mean Est. | Std. Err. | Min | Max | Std. Bias | Prop. Sig. (adj.) |
| *Habituation Effects* |  |  |  |  |  |  |  |  |  |  |  |  |
| $\rho_{growth}$ | .998 | .000 | .997 | .998 | | | .379 | .015 | .333 | .426 | | |
| *VIF* | 201.6 | 10.6 | 169.7 | 234.9 |  |  | 1.17 | .015 | 1.12 | 1.22 |  |  |
| $\gamma_{age} \text{ (mis)}$ | .001 | .025 | -.081 | .084 | **-12.17** | .021 | .233 | .022 | .1169 | .302 | **-3.02** | 1 |
| $\sigma \text{ (mis)}$ | 1.00 | .028 | .905 | 1.08 | .049 | | 1.03 | .033 | .913 | 1.12 | **.795** | |
| $\sqrt{\tau_{00}} \text{ (mis)}$ | .493 | .053 | .281 | .660 | -.124 | | .494 | .062 | .265 | .665 | -.101 | |
| $\gamma_{age}$ | .312 | .291 | -.809 | 1.56 | .030 | .109 (.065) | .299 | .025 | .233 | .369 | -.040 | 1 (1) |
| $\gamma_{prac}$ | -.312 | .391 | -1.54 | .791 | -.030 | .112 (.061) | -.299 | .051 | -.464 | -.115 | .024 | 1 (.999) |
| $\sigma$ | 1.00 | .028 | .905 | 1.08 | .031 | | 1.00 | .032 | .887 | 1.09 | -.009 | |
| $\sqrt{\tau_{00}}$ | .493 | .053 | .279 | .664 | -.124 | | .495 | .057 | .285 | .652 | -.095 | |
| *SNR Effects* |  |  |  |  |  |  |  |  |  |  |  |  |
| $\rho_{growth}$ | .998 | .000 | .997 | .998 | | | .379 | .014 | .338 | .423 | | |
| *VIF* | 202.4 | 10.3 | 169.4 | 237.6 |  |  | 1.17 | .015 | 1.13 | 1.22 |  |  |
| $\gamma_{age} \text{ (mis)}$ | -.001 | .205 | -.080 | .090 | **11.51** | .023 | -.233 | .023 | -.296 | -.149 | **2.92** | 1 |
| $\sigma \text{ (mis)}$ | .999 | .029 | .886 | 1.08 | -.028 | | 1.03 | .035 | .929 | 1.13 | **.734** | |
| $\sqrt{\tau_{00}} \text{ (mis)}$ | .498 | .055 | .294 | .709 | -.031 | | .498 | .061 | .292 | .657 | -.039 | |
| $\gamma_{age}$ | -.300 | .377 | -1.53 | .913 | .000 | .107 (.064) | -.300 | .024 | -.378 | -.222 | -.016 | 1 (1) |
| $\gamma_{prac}$ | .300 | .377 | -.917 | .155 | .001 | .104 (.062) | .301 | .052 | .150 | .492 | .022 | 1 (1) |
| $\sigma$ | .999 | .029 | .887 | 1.08 | -.041 | | .998 | .033 | .907 | 1.09 | -.047 | |
| $\sqrt{\tau_{00}}$ | .498 | .055 | .294 | .708 | -.032 | | .498 | .057 | .317 | .646 | -.027 | |

Note: Mean Est. is the expected parameter value across replicants and the Std. Err is the standard deviation of the sampling distribution. The proportion significant (Prop. Sig) represents the number of inferential tests that return a significant result for each parameter (unadjusted first, adjusted in parentheses). Std. Bias is the standardized bias estimate; values > .25 are bolded and represent unacceptable bias. The proportion significant only includes significant results where the inference is correct (e.g., significant negative effects are not included if the population parameter is positive). Rhos ( $\rho_{growth}$ ) represent the correlation between growth predictors, VIF represents variance inflation factor, gammas ( $\gamma$ ) represent the fixed effects, sigma ( $\sigma$ ) is the square-root of the residual random variance, and tau ( $\tau_{00}$ ) is the person-level random variance (square-root shown for consistency). The (mis) denotes parameters from the mis-specified (i.e., age-only) model.

#### Signal-to-Noise Ratio Effects

Another potential case of interference might be that exposure tends to increase the level of a measure, while developmental changes drive a decrease. Here the case is most clear for signals in functional neuroimaging. Individuals often feel extreme discomfort in the scanner, however, subsequent visits reduce this discomfort (Chapman, Bernier, & Rusak, 2010; De Bie et al., 2010) and lead to greater neural signal in comparison with noise (e.g., reducing motion, dual-task effects). This would then work to counteract developmental decreases we might see in a region of interest. Another example could take the form of a positive practice effect that compensates for developmental declines in performance (e.g., in old age; Koppara et al., 2015; Zimmermann & Meier, 2006). I simulated data to have a negative effect of age (*γ*_*age*_ = −0.3) but a positive effect of exposure (or practice; *γ*_*exp*_ = 0.3; Figure S3). As expected, results mirror the effects of habituation, but are reversed with positive bias in the age-only model (Table 3). As before, only when the MLMG model is paired with the appropriately-sampled data (i.e., in an accelerated longitudinal design) were stable, unbiased estimates derived for all effects in the data-generating model with appropriate inferences.

### Scenario Set 3: When Cohort Data is Appropriate

Thus far, I have highlighted the limitations of cohort-sequential for estimating multiple growth processes (e.g., age and practice) that are highly colinear, and how accelerated designs can be leveraged to address these limitations. At the risk of undermining this relatively straightforward point, I then considered scenarios where a cohort-type design would be capable of disentangling colinear growth processes of interest. However, these scenarios specifically build on the principles of an accelerated design (i.e., planned missingness), which enables proper estimation. Consider the previous scenarios. Here, experience (i.e., repeated assessments) is consistent across individuals such that each person is measured at every level of the predictor (unplanned missingness aside). In contrast, age is distributed across individuals such that each person is only measured on a subset of the levels of the predictor. While this requires an assumption that some degree of exchangeability exists across individuals with no age overlap (i.e., age convergence; Sliwinski, Hoffman, & Hofer, 2010), it allows for a decoupling of the two predictors in the sample. For a cohort-sequential design to achieve this decoupling, these features of sampling approach need to be reversed; that is, individuals are measured at all levels of the age predictor but not the experience predictor. In this way, these designs are accelerated, but with respect to the experience variable, not age.

#### Staggered School-Based Interventions

Here I simulated an application of this idea, where individuals are measured in a cohort fashion based on age, but some intervention is applied differentially across subjects (Cook & Ware, 1983). For example, perhaps we are interested in piloting how an intervention might boost reading proficiency across early education. Due to resource limitations, it might not be possible to apply treatment to all students across the study period, and therefore, we might start the intervention randomly across measurement occasions. Here I simulated a positive linear effect of both age and the intervention on reading proficiency (*γ′s* = 0.3), but a decelerating interaction effect (*γ′s* = −0.1), suggesting that intervening later in school is less effective. Previously I simulated cohort data to follow a lab-based assessment design where individuals are measured in tight clusters around discrete ages. However, in a school-based setting, we would expect individuals to be uniformly distributed in terms of age within grade, and I adopted this design for these simulations. I simulated a 4- and 5-wave observation condition to probe the effects of study duration on the challenge of multicollinearity (more observations should induce higher correlations between the growth processes). While thus far I have only focused on a relatively simple random variance structure (only a level 1 residual and level 2 random intercept), MLMGMs can accommodate further random effects just as well as other MLMs. To highlight this fact, I also included a random effect of age (*σ*_*age*_ = .15).

The MLMGM in this scenario recovered all parameters appropriately (i.e., unbiased and precise estimates), demonstrating the ability to have designs which are accelerated (i.e., planned missing) with respect to alternative growth processes (Figure 6). There were minimal differences between the 4- and 5-wave simulations with respect to appropriate parameter estimation (see Table 4 for full details). The correlations between age and intervention stage were slightly inflated for the 5-wave simulations (*M*_*ρ*_=.554, *SE* = .025) compared with the 4-wave design (*M*_*ρ*_=.532, *SE* = .024), and the false negative rate was larger but still well-controlled (5 wave: age = 0%, intervention = .2%, interaction = 1.7%; 4 wave: 0% for all predictors) due to greater variance inflation (5 wave: *M*_*VIF*_ = 1.45, *SE* = .059; 4 wave: *M*_*VIF*_ = 1.40, *SE* = .050).

**Figure 6.**
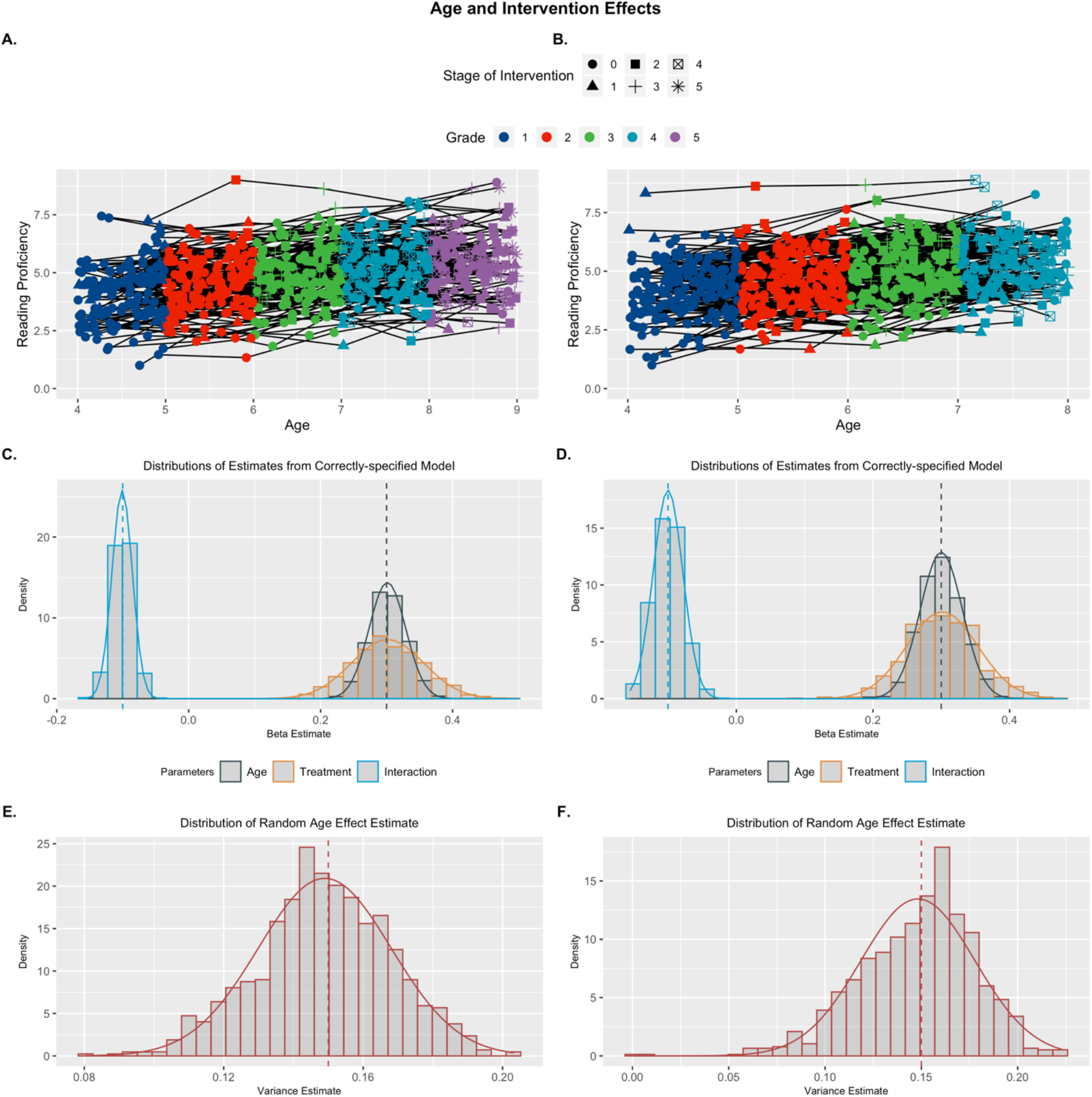
Treatment and Age Effects: Example replicants for 5-wave (A) and 4-wave (B) designs. Distributions of parameter estimates across replicants (C & D), and random effects of age (E & F) are presented for properly-specified models. Both models recovered parameters well. Generating parameters for each predictor are denoted by vertical dashed lines.

**Table 4.** Estimated Effects for Growth Models from Scenario Set 3: Cohort Models.

|  | 5-Wave Design |  |  |  |  |  | 4-Wave Design |  |  |  |  |  |
| --- | --- | --- | --- | --- | --- | --- | --- | --- | --- | --- | --- | --- |
|  | Mean Est. | Std. Err. | Min | Max | Std. Bias | Prop. Sig.(adj.) | Mean Est. | Std. Err. | Min | Max | Std. Bias | Prop. Sig.(adj.) |
| *Intervention Model* |  |  |  |  |  |  |  |  |  |  |  |  |
| $\rho_{growth}$ | .554 | .025 | .474 | .634 | | | .532 | .024 | .450 | .602 | | |
| VIF | 1.45 | .059 | 1.29 | 1.67 |  |  | 1.40 | .050 | 1.25 | 1.57 |  |  |
| $\gamma_{age}$ | .301 | .028 | .222 | .389 | .043 | 1 (1) | .300 | .031 | .186 | .406 | .003 | 1 (1) |
| $\gamma_{tx}$ | .301 | .055 | .145 | .495 | .027 | 1 (1) | .301 | .052 | .094 | .469 | .026 | .999 (.998) |
| $\gamma_{int}$ | -.101 | .015 | -.152 | -.048 | -.053 | 1 (1) | -.100 | .022 | -.156 | -.026 | -.001 | .994 (.983) |
| $\sigma$ | .501 | .017 | .444 | .556 | .034 | | .499 | .018 | .446 | .565 | -.057 | |
| $\sqrt{\tau_{00}}$ | .999 | .060 | .825 | 1.17 | -.012 | | .997 | .055 | .831 | 1.19 | -.046 | |
| $\sqrt{\tau_{11}}$ | .149 | .019 | .080 | .203 | -.053 | | .148 | .030 | .001 | .223 | -.060 | |
| *TVC Model* |  |  |  |  |  |  |  |  |  |  |  |  |
| $\rho_{growth}$ | | | | | | | .410 | .030 | .297 | .502 | | |
| VIF |  |  |  |  |  |  | 1.20 | .036 | 1.10 | 1.34 |  |  |
| $\gamma_{age}$ | | | | | | | .429 | .068 | .185 | .742 | **1.89** | 1 (1) |
| $\gamma_{stress}$ | | | | | | | .300 | .042 | .168 | .426 | -.004 | 1 (1) |
| $\gamma_{int}$ | | | | | | | -.101 | .035 | -.236 | .018 | -.026 | .829 (.700) |
| $\sigma$ | | | | | | | .500 | .019 | .444 | .557 | -.001 | |
| $\sqrt{\tau_{00}}$ | | | | | | | .999 | .055 | .810 | 1.18 | -.026 | |
| $\sqrt{\tau_{11}}$ | | | | | | | .148 | .029 | .021 | .221 | -.054 | |

Note: Mean Est. is the expected parameter value across replicants and the Std. Err is the standard deviation of the sampling distribution. The proportion significant (Prop. Sig) represents the number of inferential tests that return a significant result for each parameter (unadjusted first, adjusted in parentheses). Std. Bias is the standardized bias estimate; values > .25 are bolded and represent unacceptable bias. The proportion significant only includes significant results where the inference is correct (e.g., significant negative effects are not included of the population parameter is positive). Rhos ( $\rho_{growth}$ ) represent the correlation between growth predictors, VIF represents variance inflation factor, gammas ( $\gamma$ ) represent the fixed effects, sigma ( $\sigma$ ) is the square-root of the residual random variance, and tau ( $\tau_{00}$ ) is the person-level random variance (square-root shown for consistency). The (mis) denotes parameters from the mis-specified (i.e., age-only) model.

#### MLMGMs as a Subset of TVC Models

These results highlight the link between MLMG models and a general class of longitudinal models with time-varying covariates (TVCs). Indeed replacing intervention in the previous design with a different covariate (e.g., stress or anxiety) would be representative of a traditional single-growth process multi-level model (Curran & Bauer, 2011). For instance if we were interested in how depressive symptoms change across age during adolescence and included stress as a TVC of interest, this would follow a similar form to the MLMGMs I have demonstrated previously. However, there are some key differences between these predictors. First, the non-age growth process predictors (e.g., practice, intervention) are monotonic and intrinsically related to age (i.e., accumulating exposure does not decrease across age). While a general TVC can be correlated with age (i.e., stress also increases during adolescence), stress is not constrained to monotonic growth (e.g., stress may increase between wave 1 and 2, but decrease between 2 and 3). Additionally, the stress predictor will be subject to measurement error, whereas the MLMGM predictors I have considered thus far are truly fixed and known (e.g., number of exposure events).

I simulated data to follow this form, where adolescent depressive symptoms are modeled as a function of age, stress, and their interaction. I also included a random effect of age, mimicking the intervention scenario above. To create a better comparison with the MLMGM simulations, I simulated stress to show age-related increases, but unlike the MLMGM, there was a random error term in the model for the stress predictor (*σ*_*error*_ = .5; Figure 7B). Results showed that this error introduced bias into the estimate of the age predictor (*Bias*_*std*_ =1.89, *SE* = .068), but not the estimates for the stress or interaction predictors (compare Figure 7C with Figure 6C &D; see Table 4). Aside from this bias introduced by the measurement error in the TVC, this model performs like an MLMGM, highlighting the firm grounding of the current approach in previous methods.

**Figure 7.**
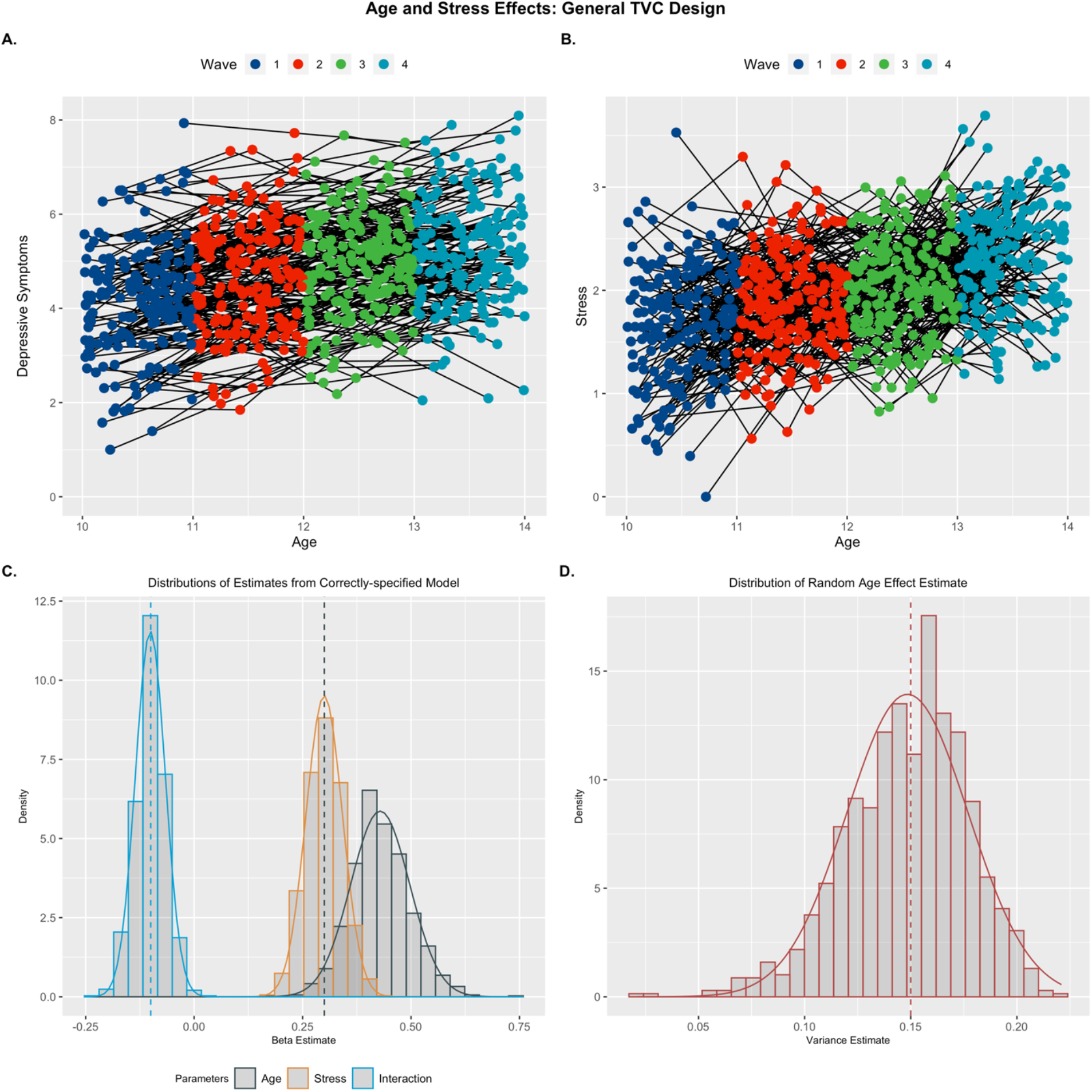
Effects from a TVC Model: Example replicant for trajectories of depression (A) and stress (B) across early adolescence. Distributions of fixed effects (C) and random age effect (D) parameter estimates are plotted. Generating parameters for each predictor are denoted by vertical dashed lines. The random measurement error in the stress predictor caused inflation in the parameter estimate of the age predictor in the model (black histogram in C).

### Scenario Set 4: Disentangling Puberty and Age Effects

The inability to jointly model the effects of pubertal hormones and chronological age has been a stumbling block for developmental science in understanding a range of phenomenon of interest (e.g., risk-taking and reward-seeking behavior). Previous work (Blakemore, Burnett, & Dahl, 2010; van Duijvenvoorde, Westhoff, de Vos, Wierenga, & Crone, 2019; Wierenga et al., 2018) has highlighted these challenges and called for implementing research designs which can address these limitations. One potential solution posed by Blakemore and colleagues (2010) was to recruit individuals at the same age, but different pubertal development status, and then follow them longitudinally. This idea fits perfectly into the framework of MLMGMs that I have presented thus far, if we think of this design as having planned missingness with respect to puberty instead of age. That is, individuals are measured across all levels of age, but not all levels of puberty. To demonstrate the ability of multi-level multi-growth models to capture these unique effects, I constructed a model to disentangle the impact of pubertal development and age on reward sensitivity across adolescence.

I considered a number of factors when constructing the simulations of age and puberty effects to better reflect the real-world nature of pubertal trends. I built ground-truth pubertal trajectories for each set of simulates from a sigmoid function (van Duijvenvoorde et al., 2019; Wierenga et al., 2018):

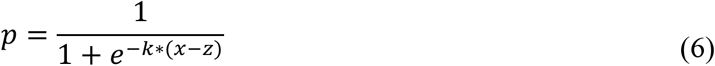

Here, *p* is the level of puberty development where 0 < *p* < 1 (lower bound = pre-puberty, upper bound = post-puberty), *z* is the mid-point of the trajectory (*p* = .5), *k* is the shape parameter which governs the steepness of the pubertal curve (steeper curves related to a shorter window of pubertal development), and *x* ranges from −4 to 4 around the mean age of the sample (*M*_*age*_ = 12 for all simulations). I drew individual values for the location parameter from a truncated normal distribution with mean 0 and bounds at −3 and 3 (*N*_*T*_[0, *σ*, −3, +3]), and values for the shape parameter from a gamma distribution (Г[10, 3]) to produce more relatively slow (> 2 years for complete transition) pubertal trajectories. I simulated a correlated multivariate distribution for the location and shape parameters such that simulates who started puberty earlier were more likely to have protracted pubertal time-courses and those who initiated puberty relatively late had faster transitions through puberty (Marti-Henneberg & Vizmanos, 1997). Examples of underlying pubertal trajectories for each condition are shown below (Figure 8A.1 – A.4).

**Figure 8.**
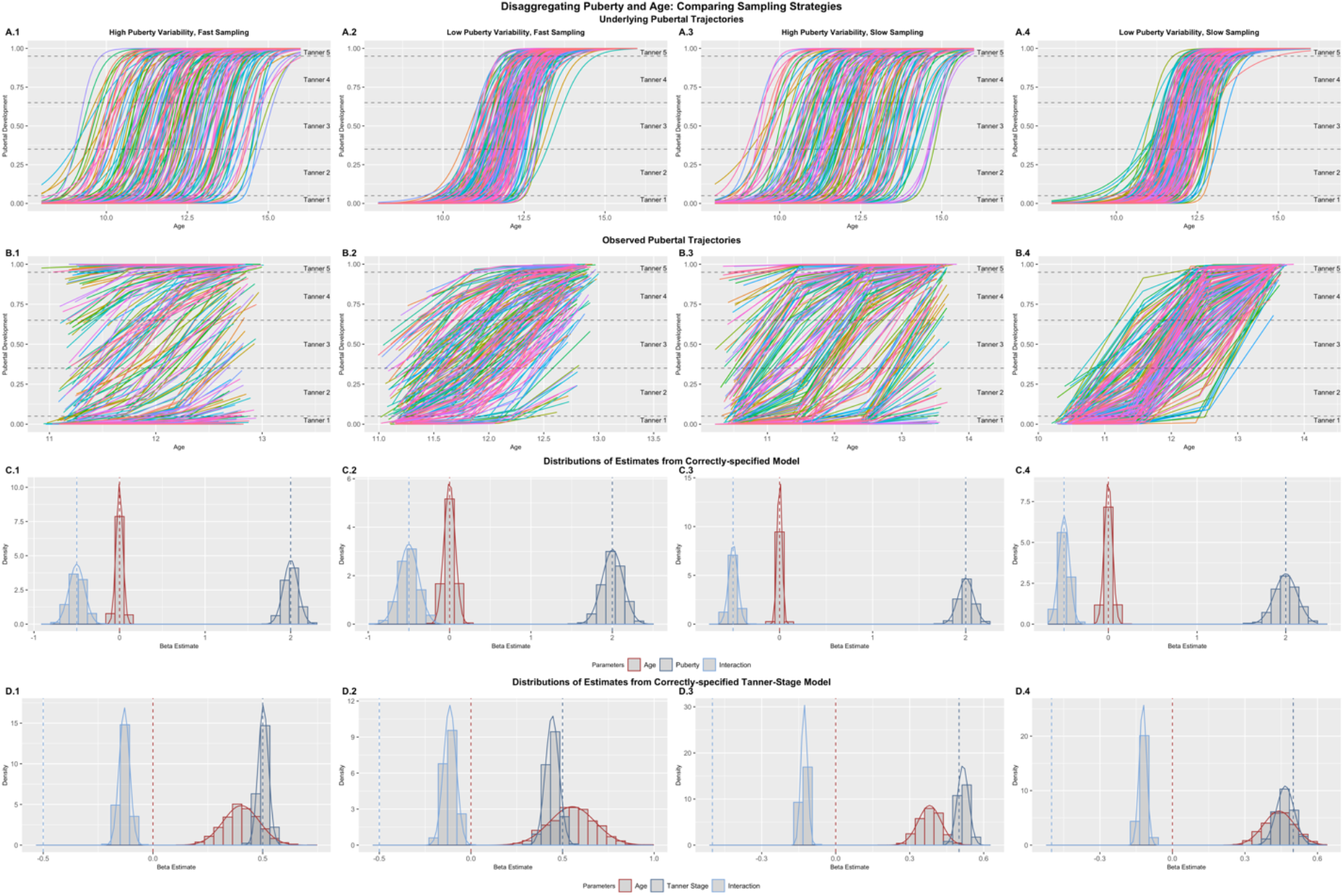
Effects from the Puberty Models: Example replicant for underlying (A) and observed (B) trajectories of pubertal development. Distributions of fixed effects are plotted for models where the puberty predictor is sampled from the continuous puberty curves (C) and models where puberty is assessed by the transformed Tanner Stage values. This tranformation introduced substantial bias into the age predictor estimates, whereas no bias was introduced when using the underlying values. Only data from the cohort-sequential model are shown.

I explored a number of variations in order to assess how MLMGMs might be used to disentangle the effects of puberty. In addition to the differences between cohort and accelerated (with respect to age) designs, I also varied the degree of variability in the midpoints of pubertal trajectories and the rate of sampling. For conditions with high pubertal variability, I set *σ* = 1.5 when drawing location parameters, and *σ* = .5 for low variability conditions. I defined high-sampling as an observation each 6 months, and low sampling as the traditional annual observation design. By centering both sampling designs around a mean of 12, this condition highlights the influence of the tails of puberty trajectories (where little change is occurring) on the ability to accurately estimate parameters of interest. Finally, while sampling from the underlying pubertal trajectories (e.g., hormonal assays) would be ideal from a measurement perspective, pubertal research in practice often relies on much coarser measurement categories (e.g., Tanner Stages; Marshall & Tanner, 1969, 1970). To reflect this reality, I defined cut-off values to transform the true pubertal measures into 5 stages. Values less than .05 reflected Tanner Stage 1 (pre-puberty), values .05 ≤ *p* < .35 reflected Tanner Stage 2 (early puberty), values .35 ≤ *p* < .65 reflected Tanner Stage 3 (middle puberty), values .65 ≤ *p* < .95 reflected Tanner Stage 4 (late puberty), and values greater than .95 reflected Tanner Stage 5 (post-puberty). I fit model with both the true and transformed Tanner scores to assess the impact of this transformation on parameter recovery. For all models, I simulated reward sensitivity to have no main effect of age (*γ*_*age*_ = 0), a strong effect of puberty (*γ*_*puberty*_ = 2), and a negative interaction effect (*γ*_*int*_ = −0.5) such that the impact of puberty on reward sensitivity is blunted at older ages. It is important to note that while the relationship between age and puberty is clearly non-linear (indeed I generated values from a truly non-linear model), the relationship between puberty and the outcome is linear, allowing for these effects to be modeled as usual in a linear model framework.

Results confirmed that the MLMGM worked to disaggregate the effects of puberty and age with high fidelity when the design was cohort-sequential with respect to age and accelerated with respect to puberty. A high sampling rate (i.e., biannual observations), coupled with a high degree of variability in pubertal timing (Figure 8B.1) showed the greatest ability to de-couple the age and puberty predictors, with between-predictor correlations near those seen in the age-practice scenarios (*M*_*ρ*_=.345, *SE* = .024). A slower sampling approach (i.e., annual observations) inflated these correlations (Figure 8B.3; *M*_*ρ*_=.599, *SE* = .025), however, with sufficient power, these effects can still be estimated reliably (i.e., false negative rates were low at the simulated effect sizes). Low pubertal variability (Figure 8B.2 & B.4) presented more significant challenges, although the high (*M*_*ρ*_=.750, *SE* = .019) versus low (*M*_*ρ*_=.898, *SE* = .007) sampling strategy was still relatively more able to attenuate the inter-predictor correlation. As in previous simulations, the high correlations do not introduce bias into the parameter estimates, but rather inflate standard errors leading to increases in false positives (for full details, see Tables 5–8). Across most conditions, an accelerated design (with respect to age) increased the correlations between age and puberty predictors relative to the respective cohort design, with the exception of the case where there is low pubertal variability and slow sampling. For conciseness, I displayed pubertal curves and parameter recovery for the cohort design results only (Figure 8); however, full details for both model types are provided elsewhere (see Tables 5–8; Figure S4-S7).

**Table 5.** Estimated Effects for Puberty Models: High Variability, Fast Sampling.

|  | Cohort-Sequential Design |  |  |  |  |  | Accelerated Design |  |  |  |  |  |
| --- | --- | --- | --- | --- | --- | --- | --- | --- | --- | --- | --- | --- |
|  | Mean Est. | Std. Err. | Min | Max | Std. Bias | Prop. Sig.(adj.) | Mean Est. | Std. Err. | Min | Max | Std. Bias | Prop. Sig.(adj.) |
| *Puberty Model* |  |  |  |  |  |  |  |  |  |  |  |  |
| $\rho_{growth}$ | .345 | .024 | .272 | .423 | | | .548 | .037 | .419 | .653 | | |
| VIF | 1.14 | .022 | 1.08 | 1.22 |  |  | 1.44 | .084 | 1.21 | 1.74 |  |  |
| $\gamma_{age (mis)}$ | .500 | .045 | .364 | .644 | **11.13** | 1 (1) | .432 | .035 | .308 | .548 | **12.41** | 1 (1) |
| $\gamma_{age^2 (mis)}$ | -.116 | .075 | -.382 | .152 | **5.14** | .287 (.196) | -.088 | .022 | -.155 | -.024 | **18.74** | .980 (.970) |
| $\sigma (mis)$ | .592 | .017 | .543 | .660 | **5.30** | | .564 | .018 | .507 | .619 | **3.48** | |
| $\sqrt{\tau_{00}} (mis)$ | .869 | .042 | .727 | 1.00 | **8.80** | | .836 | .043 | .684 | .980 | **7.87** | |
| $\gamma_{age}$ | -.001 | .039 | -.127 | .121 | -.021 | .051 (.013) | -.001 | .035 | -.121 | .108 | -.039 | .060 (.024) |
| $\gamma_{puberty}$ | 2.00 | .085 | 1.71 | 2.30 | .017 | 1 (1) | 2.00 | .098 | 1.71 | 2.31 | -.003 | 1 (1) |
| $\gamma_{int}$ | -.500 | .091 | -.814 | -.235 | -.005 | 1 (1) | -.498 | .067 | -.710 | -.249 | .036 | 1 (1) |
| $\sigma$ | .500 | .015 | .458 | .545 | -.026 | | .500 | .016 | .451 | .551 | .012 | |
| $\sqrt{\tau_{00}}$ | .497 | .033 | .396 | .602 | -.087 | | .497 | .031 | .387 | .601 | -.084 | |
| *Tanner-Stage Model* |  |  |  |  |  |  |  |  |  |  |  |  |
| $\rho_{growth}$ | .350 | .023 | .284 | .427 | | | .556 | .036 | .438 | .665 | | |
| $\gamma_{age}$ | .402 | .082 | .148 | .736 | **4.90** | .999 (.996) | .387 | .064 | .177 | .573 | **6.04** | 1 (1) |
| $\gamma_{tanner}$ | .503 | .023 | .418 | .577 | .117 | 1 (1) | .498 | .026 | .420 | .572 | -.087 | 1 (1) |
| $\gamma_{int}$ | -.130 | .024 | -.215 | -.054 | -.225 | 1 (.999) | -.125 | .018 | -.181 | -.054 | -.023 | 1 (1) |
| $\sigma$ | .500 | .015 | .458 | .545 | -.026 | | .500 | .016 | .451 | .551 | .012 | |
| $\sqrt{\tau_{00}}$ | .497 | .033 | .396 | .602 | -.087 | | .497 | .031 | .387 | .601 | -.084 | |

**Table 6.**
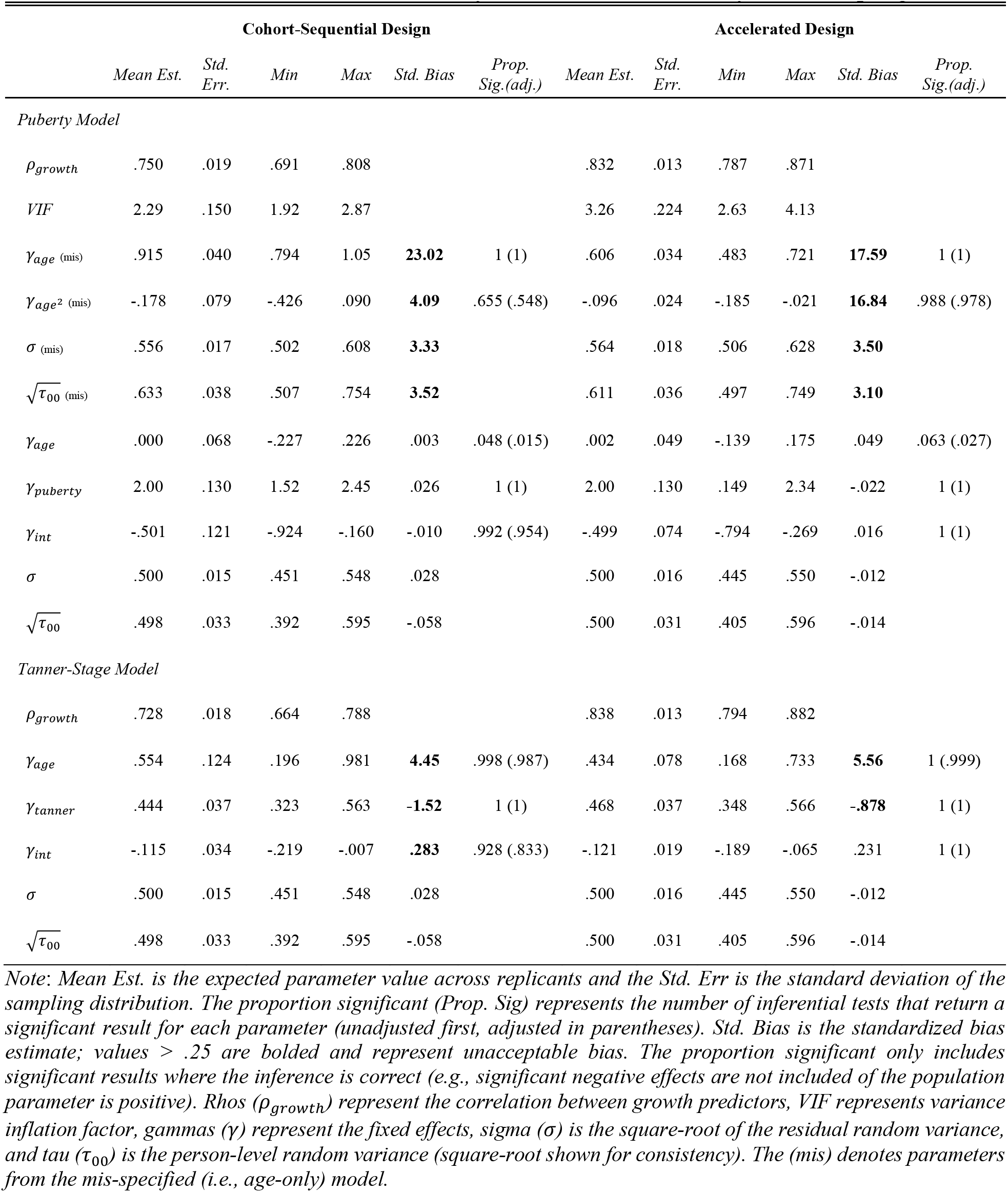
Estimated Effects for Puberty Models: Low Variability, Fast Sampling.

**Table 7.** Estimated Effects for Puberty Models: High Variability, Slow Sampling.

|  | Cohort-Sequential Design |  |  |  |  |  | Accelerated Design |  |  |  |  |  |
| --- | --- | --- | --- | --- | --- | --- | --- | --- | --- | --- | --- | --- |
|  | Mean Est. | Std. Err. | Min | Max | Std. Bias | Prop. Sig.(adj.) | Mean Est. | Std. Err. | Min | Max | Std. Bias | Prop. Sig.(adj.) |
| *Puberty Model* |  |  |  |  |  |  |  |  |  |  |  |  |
| $\rho_{growth}$ | .599 | .025 | .521 | .679 | | | .615 | .025 | .502 | .701 | | |
| *VIF* | 1.57 | .073 | 1.38 | 1.86 |  |  | 1.61 | .079 | 1.34 | 1.97 |  |  |
| $\gamma_{age \text{ (mis)}}$ | .456 | .025 | .382 | .537 | **18.50** | 1 (1) | .415 | .026 | .329 | .486 | **16.01** | 1 (1) |
| $\gamma_{age^2 \text{ (mis)}}$ | -.106 | .024 | -.176 | -.037 | **16.72** | .995 (.987) | -.084 | .014 | -.129 | -.041 | **29.96** | 1 (1) |
| $\sigma \text{ (mis)}$ | .666 | .020 | .606 | .730 | **8.12** | | .658 | .020 | .590 | .702 | **7.56** | |
| $\sqrt{\tau_{00} \text{ (mis)}}$ | .736 | .042 | .609 | .873 | **5.64** | | .744 | .041 | .604 | .889 | **5.98** | |
| $\gamma_{age}$ | .000 | .025 | -.083 | .072 | -.009 | .056 (.018) | .000 | .026 | -.084 | .084 | .001 | .065 (.023) |
| $\gamma_{puberty}$ | 2.00 | .084 | 1.71 | 2.26 | .027 | 1 (1) | 2.00 | .085 | 1.73 | 2.28 | -.031 | 1 (1) |
| $\gamma_{int}$ | -.498 | .048 | -.660 | -.337 | .042 | 1 (1) | -.501 | .045 | -.640 | -.346 | -.028 | 1 (1) |
| $\sigma$ | .499 | .015 | .456 | .557 | -.039 | | .499 | .016 | .447 | .546 | -.076 | |
| $\sqrt{\tau_{00}}$ | .497 | .032 | .387 | .589 | -.082 | | .501 | .031 | .402 | .613 | .046 | |
| *Tanner-Stage Model* |  |  |  |  |  |  |  |  |  |  |  |  |
| $\rho_{growth}$ | .606 | .024 | .534 | .677 | | | .623 | .025 | .510 | .707 | | |
| $\gamma_{age}$ | .381 | .046 | .207 | .546 | **8.20** | 1 (1) | .382 | .044 | .233 | .521 | **8.76** | 1 (1) |
| $\gamma_{tanner}$ | .516 | .023 | .439 | .600 | **.689** | 1 (1) | .512 | .023 | .447 | .586 | **.505** | 1 (1) |
| $\gamma_{int}$ | -.128 | .013 | -.169 | -.080 | -.205 | 1 (1) | -.127 | .012 | -.167 | -.092 | -.211 | 1 (1) |
| $\sigma$ | .499 | .015 | .456 | .557 | -.039 | | .499 | .016 | .447 | .546 | -.076 | |
| $\sqrt{\tau_{00}}$ | .497 | .032 | .387 | .589 | -.082 | | .501 | .031 | .402 | .613 | .046 | |

**Table 8.** Estimated Effects for Puberty Models: Low Variability, Slow Sampling.

|  | Cohort-Sequential Design |  |  |  |  |  | Accelerated Design |  |  |  |  |  |
| --- | --- | --- | --- | --- | --- | --- | --- | --- | --- | --- | --- | --- |
|  | Mean Est. | Std. Err. | Min | Max | Std. Bias | Prop. Sig.(adj.) | Mean Est. | Std. Err. | Min | Max | Std. Bias | Prop. Sig.(adj.) |
| *Puberty Model* |  |  |  |  |  |  |  |  |  |  |  |  |
| $\rho_{growth}$ | .898 | .007 | .875 | .917 | | | .853 | .007 | .829 | .876 | | |
| *VIF* | 5.21 | .334 | 4.28 | 6.30 |  |  | 3.67 | .160 | 3.19 | 4.29 |  |  |
| $\gamma_{age \text{ (mis)}}$ | .656 | .018 | .598 | .710 | **37.49** | 1 (1) | .552 | .025 | .453 | .625 | **21.72** | 1 (1) |
| $\gamma_{age^2 \text{ (mis)}}$ | -.132 | .022 | -.201 | -.048 | **16.52** | 1 (1) | -.087 | .017 | -.146 | -.041 | **24.59** | 1 (1) |
| $\sigma \text{ (mis)}$ | .586 | .018 | .526 | .638 | **4.89** | | .633 | .019 | .570 | .704 | **7.09** | |
| $\sqrt{\tau_{00} \text{ (mis)}}$ | .535 | .037 | .414 | .649 | **.943** | | .533 | .037 | .430 | .656 | **.900** | |
| $\gamma_{age}$ | .000 | .045 | -.122 | .176 | -.004 | .046 (.018) | -.001 | .036 | -.101 | .123 | -.023 | .053 (.018) |
| $\gamma_{puberty}$ | 2.00 | .129 | 1.56 | 2.36 | .002 | 1 (1) | 2.00 | .107 | 1.62 | 2.33 | .024 | 1 (1) |
| $\gamma_{int}$ | -.499 | .060 | -.680 | -.284 | .014 | 1 (1) | -.500 | .050 | -.663 | -.342 | .005 | 1 (1) |
| $\sigma$ | .499 | .015 | .457 | .555 | -.039 | | .500 | .016 | .442 | .549 | -.005 | |
| $\sqrt{\tau_{00}}$ | .497 | .033 | .398 | .606 | -.089 | | .498 | .032 | .399 | .609 | -.060 | |
| *Tanner-Stage Model* |  |  |  |  |  |  |  |  |  |  |  |  |
| $\rho_{growth}$ | .897 | .008 | .870 | .920 | | | .861 | .008 | .836 | .884 | | |
| $\gamma_{age}$ | .441 | .064 | .239 | .622 | **6.91** | 1 (1) | .383 | .053 | .228 | .554 | **7.28** | 1 (1) |
| $\gamma_{tanner}$ | .467 | .037 | .353 | .591 | **-.904** | 1 (1) | .510 | .030 | .406 | .591 | **.323** | 1 (1) |
| $\gamma_{int}$ | -.122 | .015 | -.171 | -.064 | .218 | 1 (1) | -.123 | .013 | -.166 | -.075 | .172 | 1 (1) |
| $\sigma$ | .499 | .015 | .457 | .555 | -.039 | | .500 | .016 | .442 | .549 | -.005 | |
| $\sqrt{\tau_{00}}$ | .497 | .033 | .398 | .606 | -.089 | | .498 | .032 | .399 | .609 | -.060 | |

The models with pubertal values sampled from the true underlying trajectories (Figure 8A.1-A.4) showed unbiased parameter estimates for both the main effects and the interaction term (Figure 8C.1-C.4). In contrast, coarsening the continuous trajectories into an ordinal Tanner Stage predictor introduced some bias into pubertal parameter estimates (Figure 8D.1-D.4), except in the high variability, high sampling condition (*Bias*_*std*_ = −.087, *SE* = .023). Interestingly this bias was not uniformly in one direction across all conditions. However, the bias introduced to the estimate of the Tanner Stage effect was minor in comparison to the bias introduced into the age parameter estimates (*Bias*_*std*_ *range* = 4.45 – 8.20). The biases for the interaction estimates (*Bias*_*std*_ *range* = −.210 – .283) were marginally problematic as well. Overall, the same principles that lead to successful model performance in previous scenarios held here by considering data to be accelerated with respect to puberty. However, differences due to sampling time and measurement error highlight additional challenges that will need to be overcome in real-world applications of these models to disentangle effects of puberty and age. As a general comment, models fit to the accelerated data showed inferior performance in the Tanner Stage models, with higher inter-growth process correlations without improving estimate biases.

## Discussion

A key challenge for the study of change over time is disentangling effects of correlated predictors across time. While this challenge is not unique to longitudinal designs, they are uniquely problematic since predictors like age, experience, and puberty monotonically increase across time. Even if the true effects of these predictors are nonlinear (e.g., quadratic), current modeling approaches necessitate that linear terms also be included. When these linear terms are too highly colinear, they can introduce significant problems for proper estimation of model parameters and impact the substantive conclusions about inflection points and sensitive periods in developmental trajectories. However, simply excluding one of the correlated predictors threatens the internal validity of the model and introduces significant bias into the retained parameters except under very restrictive assumptions (i.e., the true effect of the excluded predictor is zero). Fortunately, leveraging a natural extension of linear mixed-effects growth models in combination with appropriate sampling designs offers a solution to these challenges. Here I outlined a multi-level, multi-growth model (MLMGM) form and highlighted how it can be flexibly used across a number of scenarios to address questions from substantive developmental theory.

### Quantifying the Effects of Repeated Exposure

The strength of repeated-measures designs (i.e., the ability to assess within-person change), can simultaneously be a threat to internal validity. That is, individuals may change in their response to an instrument (e.g., survey, fMRI task) due to being assessed multiple times independent of changes that are due to developmental effects (Bell, 1953; Palmore, 1978; Telzer et al., 2018). Whether because of familiarity or practice with a task, changing sensitivity of certain questionnaire items (e.g., drug use), or increased comfort with the testing environment, the effect of repeated exposure can introduce bias into estimates of developmental trajectories. I began by outlining a number of scenarios where this bias is highlighted. Except in scenarios where the true of effect of repeated exposure was zero, the presence of these effects posed challenges at multiple stages of model estimation and interpretation. However, merely including relevant predictors does not necessary ameliorate all of these challenges. Traditional cohort-longitudinal designs, where all individuals are assessed at every age levels (Figure 1B), limit the utility of including predictors of repeated exposure because the high correlation between age and assessment (*M*_*ρ*_ ≈ .998) limits the ability to uniquely model these effects with any precision, or detect significant effects when they do exist (i.e., standard errors are highly inflated). I showed that only MLMGMs combined with accelerated-longitudinal designs could reliably recover the underlying model generating parameters under commonly-encountered conditions. I highlight how this approach can address several specific challenges.

#### Bias from Omitted Predictors

Across all of the presented scenarios, I first fit models that follow conventional practices (Curran et al., 2010; Duncan, Duncan, & Hops, 1996; Herting et al., 2018; Sliwinski et al., 2010; Soden et al., 2015; Telzer et al., 2018), where only the effects of age are assessed as a growth process in the model (i.e., single-growth models). These models highlighted the various challenges that can arise from omitting relevant predictors. In cohort-sequential data, the effect of repeated measures (e.g., practice, habituation) was almost completely absorbed into the parameter estimates for age due to the high correlations between predictors (Frees, 2001; Kim & Frees, 2006). Depending on the nature of the two effects, this lead to serious over- or under-estimation of the true age effect. Here we can see one of the advantages of the accelerated-longitudinal design even in the mis-specified model. While the effects of age are still biased, the magnitude of this bias is substantially lower than in the cohort data (standardized values of ~ 3 versus ~ 11.5) due to the attenuation of the correlations between predictors (*M*_*ρ*_ ≈ .380). However, this attenuation allows for even greater validity in modeling developmental trajectories, since both age and the repeated measure predictor can be included. When including these multiple growth predictors, the model appropriately estimates the independent effects of age and repeated exposure from accelerated data. While effects are unbiased in aggregate with multiple predictors in the cohort data, the parameters from any one replicant (i.e., study) may be quite biased due to instability in the estimates, and the likelihood for false-negatives is high (> 85 %) due to inflated standard errors (> 14 times as large as if predictors were uncorrelated).

Notwithstanding the issues highlighted in these scenarios, it is likely that there are scenarios where there are truly no (or at least minimal) effects of repeated exposure on an outcome of interest. Under these conditions, the MLMGM with accelerated data would perform well (see age effects only simulations), but would be a less-ideal approach due to the reductions in power in these designs relative to cohort-sequential studies. However, in real data contexts, there is no way to test this assumption, while the MLMGM with accelerated data makes this a specific, testable hypothesis without compromising the estimation of the age effect. At high sample sizes, the relatively minor tradeoff in power makes MLMGMs with accelerated designs an attractive way to assess developmental trajectories while controlling for confounds due to the effect(s) of repeated assessment. As demonstrated by the simulation results, failure to account for these omitted effects could lead to biased estimates and contribute to a lack of reproducibility and generalization of effects across studies if the effects of repeated exposure vary between measures, populations, or other assessment parameters.

#### Bias in Models with Higher-Order Effects

The issue of high correlations between predictors impacts linearly-related independent variables, however, there are any number of situations where the relationship between growth predictors and outcomes of interest might be expected to be non-linear, or some combination of linear and non-linear effects. Because of the centering method for creating product terms (i.e., interactions or polynomials; Aiken et al., 1991), the predictors for these higher order effects will be uncorrelated with the linear terms in the model. The simulations with a quadratic effect of age highlight this fact, as this term is estimated without bias and with high precision in all models, including the ones with cohort sequential data (Figure 4). Unfortunately, this does not address the issue of correlated linear predictors, which are necessary to properly model any higher-order effects, nor does it solve issues for model-implied features of the developmental trajectories. These implied features can include the rate of the linear trend, or where the quadratic or interaction inflection (i.e., vertex) occurs within the overall developmental trajectory. Because of the instability in the linear term, the same properly specified quadratic effect can lead to implied inflections that vary widely. For instance, the implied vertex in the mis-specified (i.e., age-only) model using a cohort-sequential design ranged across 5 years (15-20), and in the properly-specified model, the implied vertex of the effect varied across more than a full decade. Because real studies do not have the benefit of obtaining a sampling distribution, results from any one study could lead to radically different substantive conclusions based on parameter instability. This complication could further decrease replicability of effects across samples, even though the point estimates of product term effects could be nearly identical.

### Generalizing the Idea of Accelerated Designs

While the distinction between experience- (e.g., practice, habituation, etc.) and age-related changes in outcomes of interest is a persistent issue, the utility of the MLMGM is not limited to these effect. Experience effects can often be thought of as a nuisance variable (although see McCormick et al., *preprint* for an exception) that contaminates the developmental effect. As such, MLMGMs can be used to partial out variance associated with repeated assessment. However, in other contexts, both growth processes may be of interest and the focus can be on characterizing trajectories across both levels of time. In many of these scenarios, it might be that accelerated designs (as typically conceived) are impossible or undesirable. Nevertheless, the MLMGM framework can be leveraged. The key insight to understand how MLMGMs separate different growth effects; by measuring all individuals across the full range of one growth predictor, while measuring individuals on only a subset of the levels of the other predictor. In the age/experience simulations, every individual is measured at three occasions (i.e., across all levels of the experience predictor) but on only a subset of ages. This fits perfectly within a generalized framework of planned missingness in study design (Little & Rhemtulla, 2013; Rhemtulla & Little, 2012). Under this broad framework, there is nothing (from a modeling standpoint; design feasibility may impose some limits) that prevents researchers from flipping that idea for other questions about multiple growth processes.

I presented one such example, where individuals are measured at all levels of age (within the scope of the study), which results in a cohort-sequential data frame, but where the levels of the intervention vary across individuals at any given timepoint. I also showed how the MLMGM is a special case of a general growth model with a time-varying covariate (TVC) with a model distinguishing effects of age and stress on depression (Curran & Bauer, 2011). However, the multi-growth model introduces challenges that may not impact more general TVC models. TVCs, like stress, can fluctuate non-monotonically across measurement occasions, even when there is a general trend across age (e.g., stress generally increases across the transition to adolescence). In contrast, the predictors in growth-processes (including age, experience, and puberty) cannot decrease from one measurement occasion to the other, which drives the need to leverage accelerated designs to attenuate otherwise near-perfect multi-collinearity. The advantage of these multiple-growth predictors, relative to other TVCs, is that there can potentially be very low – or non-existent – measurement error (although this may not always be the case) since the number of assessments or interventions for each individual is truly fixed and known. This set of scenarios, where planned missingness occurs in a growth predictor other than age is the richest target for novel and innovative future work and where existing cohort-sequential datasets can be used to empirically test the types of effects I have presented here.

### Using Multi-Level, Multi-Growth Models to Address Developmental Theory

Finally, I applied the principles derived from the first three sets of scenarios to demonstrate an approach for disentangling the effects of pubertal and age-related development. This challenge is a long-standing one in the developmental literature (Blakemore et al., 2010; van Duijvenvoorde et al., 2019), and interestingly, results suggest that data needed to address this gap likely already exist. In particular, I showed that cohort-sequential designs could be leveraged to estimate both age and puberty effects with reasonably low correlations, especially when sampling occurs more often than typical annual visits (*M*_*ρ*_ = .345). However, even though annual observations increased this correlation (*M*_*ρ*_ = .599), a sufficiently powered design can overcome these challenges (the current simulations had 188 individuals measured four times) with standard errors being only ~1.25 times larger than expected. In the universe of longitudinal studies (a strong conditional statement), this kind of design exists or will with increasing frequency in the coming years (e.g., ABCD study) for a range of theoretical questions. However, the limiting factor here appeared to be the variability of pubertal trajectories within the observation period, a fact which is incredibly difficult to account for at the onset of data collection. In simulations with low pubertal variability, the correlations between age and puberty reached potentially problematic levels, especially with slower (i.e., annual) observations (*M*_*ρ*_ = .898). One strategy for addressing this limitation could be to sub-sample individuals from a larger study to maximize the amount of pubertal variability in the sub-sample, however, this would have to be done with care to ensure that the properties of the sub-sample match the larger study characteristics, especially avoiding inadvertently selecting on the outcome (Elwert & Winship, 2014). Of course, this could be further complicated by the likely need to separate out effects by gender (something I deliberately chose to ignore in the current simulations), but in principle the same approach could be done for each group if ad hoc sampling techniques did not (by chance or design) result in high levels of pubertal variability. Alternatively, high-density sampling attenuated the correlation between predictors even with low pubertal variability, offering researchers multiple avenues for building these models.

While the MLMGM framework was able to appropriately capture pubertal and age-related effects when using measurements of the underlying trajectory, coarse categorization of those values to an ordinal scale (representing Tanner Stages) introduced bias into both the estimated effect of Tanner Stage and uniformly inflated the estimate of age. Importantly, this bias was not due to meaningful differences in the correlations between the growth predictors (see Tables 5–8). Results from the TVC simulations suggest that the measurement error introduced by transforming the measure of puberty may be absorbed into the age estimate, leading to the bias. This suggests that age effects in models using Tanner Stage as a predictor may need to be interpreted with caution, as they may be biased upwards. The bias in the Tanner Stage estimate itself was smaller and directionally inconsistent across simulation conditions (Figure 8D), but also suggests that unlike the effect of stress in the TVC scenario, the coarse categorization also impacts the estimate of the pubertal effect. These results suggest that continuous measures of puberty (e.g., hormone concentrations) are likely to be more reliable in their effect estimates, even before taking into consideration the increased measurement precision of assays versus self-report or even physician evaluation. This is true, despite the near perfect correlation between the underlying puberty measure and the Tanner Stage (*M*_*ρ*_ = .965 – .985). While continuous measures of puberty are often more expensive and can be challenging to collect, these results suggest that they are likely necessary. In studies with more-coarse measures, care should be taken in drawing overly-confident conclusions in light of the bias being introduced.

Finally, while not the main focus of the current analyses, I did assess how the use of an accelerated design (with respect to age) impacted results (see Tables 5–8 for details; Figures S4-S7). In general, these models performed similarly to those fit to cohort data and were able to deliver unbiased estimates for the models using underlying pubertal trajectories. The drawback was that these designs inflated the correlation between the puberty and age predictors, increasing standard errors for all estimates. However, it is important to note that while the kind of accelerated design I simulated reflects common practice, it could be modified to work as well or better than the cohort design presented here. For example, instead of planning the missingness around puberty, researchers could use an accelerated (i.e., missing age) design and measure individuals across the complete range of puberty instead, or sub-sample from a large accelerated study to approximate this design. This is obviously a more challenging design to realize than one where researchers can simply track age, given the variability in timing and rate of pubertal development and potentially complicated relationships between underlying maturation and easily-identifiable physical markers. Nevertheless, some theoretical questions may require individuals sampled across the full range of puberty to properly address, making a focused study of this nature extremely valuable.

### Future Work

While the current study suggests that MLMGMs offer a promising framework for simultaneously modeling growth processes that co-occur across time, there remain two avenues for future work to explore. As I have hopefully demonstrated with a small subset of potential scenarios, these models can be applied to understand a wide array of substantive theoretical questions. In addition to being useful for testing novel questions, MLMGMs could be used to re-visit previous data where simultaneous growth processes were not considered (i.e., age-only models). Additionally, the findings here highlight the clear importance of sampling design for fitting models appropriately in longitudinal data. While this is hardly a new revelation (Bell, 1953; Kraemer et al., 2000; Louis et al., 1986; Palmore, 1978; Van’t Hof et al., 1977), it bears repeating due to the crucial role design plays in reducing correlations between growth predictors and keeping standard errors appropriately narrow. With these considerations in mind, I hope that the current work can help guide analysis of current data, as well as motivate new data collection with planned missing designs to disentangle multiple growth processes that occur across time.

From a methodological perspective, there are additional steps that could broaden our knowledge about how MLMGMs perform across a variety of conditions that are less ideal, but nevertheless common. For instance, while I attempted to choose sample sizes that would reasonably reflect reality and avoid potential issues of power, the sample sizes encountered in practice likely vary widely (e.g., > 1000 might be reasonable for a school-based survey study, but 100 might be more typical of neuroimaging studies). In particular, future work should seek to characterize the boundary conditions where correlations between growth predictors are too inflated for a given sample size to yield satisfactory inferential performance. While this work can build off principles with work done with correlated predictors in general (Shieh & Fouladi, 1991), the unique characteristics of growth predictors (e.g., monotonic across time) may require innovative designs and tailored solutions to decouple them. Additionally, while the accelerated design takes advantage of planned missingness, longitudinal studies are often subject to missing data mechanisms that are more pernicious, including attrition and other non-random processes (Enders, 2011; Schafer & Graham, 2002). While I did test the effect of additional random missingness (Table S1), this missingness was completely at random and the impact of problematic forms of missingness needs to be explored to improve the utility of MLMGMs in practice. Finally, growth factors in the scenarios that I explored all varied at each measurement occasion at level 1. For instance, age or puberty was necessarily different at each observation (although the degree to which they changed might differ). However, it seems possible that growth factors might change differentially across time and potentially enter as predictors at different levels of the model. An example could arise if researchers were interested in modeling changes within-session in addition to between-session growth, where age would be consistent across all levels of the within-session observations. One potential issue is that nesting under a predictor like age or wave might cause estimation or inferential problems that are not entirely clear. Additional work will be needed to clarify this point.

## Conclusions

I proposed an natural extension of the multi-level growth model to accommodate estimating multiple growth process simultaneously: the multi-level, multi-growth model (MLMGM). To deal with the correlations between growth predictors (e.g., age and repeated exposure), I demonstrated how accelerated-longitudinal designs could attenuate these predictor relationships through planned missingness. I further generalized the idea of accelerated designs to show that you could plan missingness in growth predictors other than age (e.g., treatment) and how these models behaved similarly to general growth models with time-varying covariates. Finally, I demonstrated how MLMGMs could be used to address the thorny substantive question of simultaneously estimating the disparate impacts of puberty and age on behavior and cognition. Taken together, this work motivates innovation in developmental designs and modeling of trajectories across time across a broad domain of empirical research.

## Notes

https://mccormickneuro.github.io/publication/preprint-mlmgm/mlmgm.html

### Competing Interest Statement

The authors have declared no competing interest.

